# Chemical warfare in the rice plant rhizosphere: Diverse *Pseudomonas* antagonize the green alga *Chlamydomonas*

**DOI:** 10.64898/2026.11.11.765871

**Authors:** Xiaoqi Yang, Prateek Shetty, Bhoomika Vajrangi, Inam UI Haq, Maria Mittag

## Abstract

Algae and bacteria form dynamic partnerships that shape diverse ecosystems. The model green microalga *Chlamydomonas reinhardtii* is often found in paddy rice fields, where the diversity and functional characteristics of its bacterial partners remain largely unexplored. Here, we isolated 12 bacterial strains from the rice rhizosphere and its surrounding soil. Phylogenetic analysis based on *16S rRNA* gene sequences revealed that eight isolates belong to the genus *Pseudomonas*. Three *Pseudomonas* rhizosphere strains exhibited antagonistic effects against *C. reinhardtii*, negatively impacting its growth, pigmentation and cell morphology. Close physical associations between the three antagonistic *Pseudomonas* strains and *C*. *reinhardtii* were observed in liquid cultures. Whole genome sequencing identified the antagonistic strains as *Pseudomonas alloputida*, *Pseudomonas tohonis*, and *Pseudomonas asiatica*. The three bacterial genomes encode several factors well-known for mediating chemical warfare. Notably, *P*. *tohonis* bears a gene cluster encoding the phenolic toxin, 2,4-diacetylphloroglucinol (DAPG). DAPG inhibits the growth and alters the cell morphology of *C*. *reinhardtii*. These findings expand our understanding of the diversity of bacterial partners associated with *C*. *reinhardtii* and highlight the role of *Pseudomonas* rice rhizosphere-associated bacteria as potent green algal antagonists. They harbor a reservoir of common functional traits that negatively influence green algal population dynamics.

## 1 | INTRODUCTION

Photosynthetic eukaryotic microorganisms, called microalgae, account for approximately half of the global net primary productivity together with cyanobacteria and are of high ecological relevance for life on Earth (Field et al. 1998). They are widely distributed across marine, freshwater, and terrestrial ecosystems. Bacteria have coexisted with microalgae for billions of years in these diverse ecosystems to form intimate interactions that regulate the stability of the ecological community and influence geochemical cycles (Cirri and Pohnert 2019). Microalgae and bacteria mutually influence each other’s physiology and metabolic properties (Abate et al. 2024; Burgunter-Delamare et al. 2024). Microalgae can secrete substances to interfere with quorum sensing regulation in bacteria (Teplitski et al. 2004). Environmental abiotic factors also have a significant influence on the interaction between microalgae and bacteria (Abate et al. 2024).

*Chlamydomonas reinhardtii* is a powerful microalgal model to study algal-bacterial interactions, including mutualistic and antagonistic relationships (Calatrava et al. 2023, 2024). These interactions are mediated by the exchange of primary and secondary metabolites provided by the interacting partners (Burgunter-Delamare et al. 2024). For example, a recently evolved B_12_-dependent strain of *C*. *reinhardtii* forms a mutualism with B_12_-producing bacteria (Bunbury et al. 2022). *Mycetocola lacteus* undergoes a mutualistic partnership with *C*. *reinhardtii*, which supplies vitamins B_1_, B_3_, and B_5_, as well as methionine for bacterial growth (Carrasco et al. 2024). *Methylobacterium aquaticum* relieves the inhibitory effect of elevated auxin levels on *C*. *reinhardtii* cell multiplication and chlorophyll degradation, and its growth is mutualistically enhanced by the alga (Calatrava et al. 2024). In contrast, the soil bacterium *Pseudomonas protegens* Pf-5 antagonizes *C*. *reinhardtii* using a blend of toxins that reduce algal cell growth, alter cell morphology and even deflagellate and lyse the algal cells (Aiyar et al. 2017; Hotter et al. 2021; Rose et al. 2021).

Although complex bacterial interactions with *C. reinhardtii* have been heavily characterized *in vitro* (Calatrava et al. 2023), understanding their true ecological significance requires examining natural habitats where these organisms coexist. Originally isolated from potato field soil (Harris 1989), *C*. *reinhardtii* assembles a diverse, conserved microbiota that resembles the root microbiota of land plants (Durán et al. 2022). Beyond generic farm soils (Lingappa et al. 2026), this microalga is also identified in paddy rice soils (Barrett et al. 1982; Ghasemi et al. 2008). Organic paddy rice soils exhibit greater temporal variability in bacterial community dynamics than conventional paddy soils (Zhu et al. 2024). Root-associated microbiomes of rice have been extensively explored at the bacterial genus level, expanding our understanding about the composition of the bacterial community (Edwards et al. 2015; Dastogeer et al. 2022). It has been found that the rhizosphere niche is especially important for land plant growth and that the unique oxic-anoxic interface of rice soil accommodates a variety of microbes (Islam et al., 2024). Previous studies have demonstrated that the bacterial populations dominate the rice rhizosphere, exhibiting at least twofold greater abundance than in bulk soil (Hussain et al. 2012; Breidenbach et al. 2016). However, the bacterial diversity in the rice rhizosphere is close to, or even less than in the bulk soil (Edwards et al. 2015; Breidenbach et al. 2016). The bacterial community in the rice rhizosphere includes Proteobacteria, Acidobacteria, Actinobacteria and Chloroflexi phyla (Ding et al. 2019). In paddy rice soil, carbon sequestration is done by rhizospheric microorganisms that can directly assimilate CO_2_ through photoautotrophy (Islam et al. 2024). The composition of the rhizosphere microbiome is also shaped by root exudates, including organic acids (Bais et al. 2006). Rice soil is in general enriched in acetate (Hori et al. 2007), a readily assimilated carbon source for efficient growth of *C. reinhardtii* (Harris 1989; Perrineau et al. 2014).

Our study aimed to find natural occurring rice field bacteria at the species level with the potential to interact with *C. reinhardtii* in a positive or negative way. Genomic characterization of these bacteria shall extend our understanding about the genomic basis in algal-bacterial interaction. Thus, we isolated bacteria from the rice rhizosphere and its surrounding soil in an organic managed rice field. We used whole-genome sequencing to identify three *Pseudomonas* species, *Pseudomonas alloputida*, *Pseudomonas tohonis*, and *Pseudomonas asiatica*, that antagonize *C. reinhardtii*. The three bacterial genomes encode factors well-known for mediating chemical warfare. The genome of *P. tohonis* carries a biosynthetic gene cluster (BGC) encoding 2,4-diacetylphloroglucinol (DAPG), a polyketide produced by some *Pseudomonas* strains. DAPG can protect land plants (Suresh et al. 2022) but exerts a toxic effect against *C. reinhardtii* (Rose et al. 2021). Understanding how rhizosphere bacteria, known to protect crops, restrict microalgal populations provides essential insights into the microbial dynamics regarding nutrient cycling and planetary soil health.

## 2 | Materials and Methods

### 2.1 | Sample Collection and Bacterial Isolation

Rice soil and rhizosphere samples were collected from an organic rice field in Ottobiano, Pavia, Italy (45°09′01.9″N, 8°49′42.5″E). The field had been managed under organic farming practices for 21 years and was cultivated continuously with rice for 25 years until 2025. Rice (variety Rosa Marchetti) was cultivated under flooding irrigation. The field was managed under a minimum tillage system, with cover crop residues incorporated into the soil as fertilizer. The soil was classified as sandy, with a moist condition and a pH of 7.2. The rice plants were seeded on May 31, 2025. Field sampling was performed on June 30, 2025, corresponding to the early tillering stage of rice (BBCH 21–23). At sampling, the air temperature was 35 °C, the soil was moist, and soil temperatures were 30 °C and 27 °C at depths of 0 cm and 5 cm, respectively.

Field description, as well as soil and rhizosphere sampling were carried out with Innova-Tech (Ottobiano, Italy). Samples were taken using a shovel that had been disinfected with 70% (v/v) ethanol and allowed to air dry before sampling. For rhizosphere samples, roots were gently shaken to remove loosely adhering soil. Subsequently, roots with adhering soil were placed in a sterile 50 mL conical tube containing 25 mL sterile 1x phosphate-buffered saline (PBS; 6.89 mM K_2_HPO_4_, 2.50 mM KH_2_PO_4_, and 140 mM NaCl; pH 7.2) and vortexed for 15 s to dislodge the rhizosphere soil. An aliquot (3 mL) of the resulting rhizosphere-PBS suspension was then transferred to 15 mL Tris-Acetate-Phosphate (TAP) medium (Harris 1989). For bulk soil samples, 0.5 g of soil was taken and inoculated into 15 mL TAP medium. The cultures were enriched by incubating them under continuous light (50 μmol photons m⁻² s⁻¹) at 23 °C with orbital shaking (100 rpm). 50 µL of each sample were spread onto TAP agar plates and incubated at 23 °C for 4–5 days. Colonies exhibiting distinct morphological characteristics were selected and sequentially re-streaked onto fresh Luria-Bertani (LB, Carl Roth, Karlsruhe, Germany) agar plates for two additional rounds to obtain an axenic bacterial colony. The axenic status of the bacterial colony was confirmed by optical microscopy observations and *16S rRNA* gene sequencing. Each re-streaked culture was incubated on LB plates at 23 °C for 3–5 days until colonies developed.

### 2.2 | Strains and Culture Conditions

*C*. *reinhardtii* strain SAG 73.72 was obtained from the Culture Collection of Algae at Göttingen University (SAG). It is an isolate of Tsubo’s strain C8, which originates from 21 gr (Sager) that derives directly from the originally isolated field strain from Smith, 1945 (Harris 1989). *C*. *reinhardtii* cells were grown in acetate-containing TAP medium in a light-dark regime of 12 h light and 12 h darkness (LD 12:12) with a light intensity of 50 μmol photons m^−2^ s^−1^ and stirring (100 rpm) at 23 °C. Algal cells grow mixotrophically under these conditions.

### 2.3 | Cocultivation of *C*. *reinhardtii* and Bacteria

For cocultivation of algae and rice field bacteria on agar plates, a TAP plate (2% agar) was overlaid with a 0.5% TAP agar top layer containing 1 × 10^6^ *C*. *reinhardtii* cells mL^-1^. 50 μL of an overnight bacterial culture adjusted to an optical density at 600 nm (OD_600_) of approximately 0.3 was applied to a restricted area of the solidified plate. The plates were incubated at 23 °C in an LD12:12 cycle with a light intensity of 50 μmol photons m^−2^ s^−1^.

For cocultivation of algae and bacteria in liquid culture, *C*. *reinhardtii* (1 × 10^5^ cells mL^-1^) and bacteria (2.5 × 10^7^ cells mL^-1^) were grown in 50 mL conical flasks containing 15 mL TAP medium as described for algal monocultures. Algal cell density was determined by counting using an improved Neubauer chamber (No. T729.1, Carl Roth, Karlsruhe, Germany). Bacterial cell density was determined as colony forming units (CFU) by plating serially diluted cultures on LB agar plates followed by colony counting after two days of growth at 26 °C. Correlation curves between bacterial density and OD_600_ values were established for R-Bac-5r, R-Bac-6r, and R-Bac-8r, as shown in Figure S1. OD_600_ was measured using a Jenway 6305 spectrophotometer (Jenway, Stone, UK), with LB medium alone used as the blank.

### 2.4 | *16S rRNA* Gene Sequencing and Phylogenetic Analysis

The *16S rRNA* gene region of the 12 bacterial isolates was amplified using universal bacterial primers. The forward primer was 5′-AGAGTTTGATCCTGGCTCAG-3′ and the reverse primer was 5′-GGTTACCTTGTTACGACTT-3′. PCR amplification was performed in a total reaction volume of 25 μL, containing PCR buffer, MgCl₂, dNTPs, forward and reverse primers, Taq DNA polymerase, and bacterial genomic DNA as the template. The PCR program consisted of an initial denaturation step at 98 °C for 30 s, followed by 30 cycles of denaturation at 98 °C for 10 s, annealing at 55 °C for 30 s, and extension at 72 °C for 30 s, with a final extension step at 72 °C for 5 min. The amplified PCR products were separated by 1% agarose gel electrophoresis to verify the expected fragment size. The confirmed amplicons were purified using a commercial PCR purification kit and subsequently submitted to Eurofins Genomics (Ebersberg, Germany) for sequencing.

The *16S rRNA* gene sequences of the bacterial isolates were aligned using the MUSCLE algorithm implemented in MEGA12 software (Kumar et al. 2024). A phylogenetic tree was subsequently constructed using the Maximum Likelihood (ML) method (Felsenstein 1981) based on the best-fit nucleotide substitution model selected in MEGA12. The robustness of the phylogenetic relationships was evaluated by bootstrap analysis with 1,000 replicates. The phylogenetic analysis included the *16S rRNA* gene sequences of the isolates obtained in this study and closely related reference type strains retrieved from the NCBI GenBank database.

### 2.5 | Genome Isolation, Sequencing, Assembly and Annotation

Genomic DNA from the bacterial strains R-Bac-5r, R-Bac-6r, and R-Bac-8r was isolated for next-generation sequencing. For this purpose, a 10 mL bacterial culture was grown in LB medium at 26 °C for 16 h. Cells were harvested by centrifugation, and the resulting pellet was used for total genomic DNA extraction with the DNeasy PowerSoil Pro Kit (QIAGEN, Hilden, Germany) according to the manufacturer’s instructions. Library preparation and sequencing were performed by Eurofins Genomics (Ebersberg, Germany). Raw paired-end Illumina reads were processed to remove adapter sequences and low-quality bases using fastp (v0.24.0, Chen et al. 2018). The trimming parameters included the automatic detection of adapters for paired-end data and the removal of low-quality bases from both the 5’ and 3’ ends. Reads were filtered to ensure a minimum length of 45 bp and a minimum Phred quality score of 20. Cleaned reads were assembled *de novo* using the Unicycler (v0.5.1, Wick et al. 2017) pipeline in “normal” mode. The quality and completeness of the resulting assemblies were evaluated using CheckM2 (v1.1.0, Chklovski et al. 2023). CheckM2 utilizes a machine learning-based approach to predict genome completeness and contamination rates by comparing the assembled sequences against a reference set of lineage-specific marker genes. Initial taxonomic classification was performed using the Type (Strain) Genome Server (TYGS, Meier-Kolthoff and Göker 2019). To refine the identification and for isolates that could not be initially identified through the TYGS database, the Genome Taxonomy Database Toolkit (GTDB-Tk, Chaumeil et al. 2019) classify_wf workflow was employed to provide species-level assignments. Following GTDB-Tk identification, additional reference genomes of *Pseudomonas alloputida* NMI24_14 (GenBank Accession: GCA_021283545.2) and *P*. *protegens* Pf-5 (GenBank Accession: GCA_000012265.1) not originally present in the TYGS database were incorporated into a secondary TYGS analysis to generate a comprehensive genome-scale phylogeny. Pairwise intergenomic distances were calculated using the Genome BLAST Distance Phylogeny (GBDP) approach under the ’trimming’ algorithm and formula d5. Digital DNA-DNA hybridization (dDDH) values and confidence intervals were calculated using Genome-to-Genome Distance Calculator (GGDC 4.0). Average nucleotide identity (ANI) analysis was performed using the integrated prokaryotes genome and pan-genome analysis web services (IPGA) v1.09 (Liu et al. 2022). Phylogenetic trees were inferred using FASTME 2.1.6.1 from the resulting intergenomic distances. The final genome-scale phylogenetic tree was visualized and plotted using the Interactive Tree of Life (iTOL) webserver (Letunic and Bork 2024).

### 2.6 | Comparative KEGG Functional Annotation of *Pseudomonas* spp. Genomes

All four *Pseudomonas* spp. genomes were subjected to genome annotation using the same RAST annotation pipeline to minimize annotation-related differences among genomes (Overbeek et al. 2014). The predicted protein sequences of four *Pseudomonas* spp. genomes were submitted to the KEGG Automatic Annotation Server (KAAS) for functional annotation (Moriya et al. 2007). The resulting KEGG Orthology (KO) assignments were downloaded and used for all subsequent comparative analyses. A KO presence–absence matrix was generated by integrating KO annotations across all four genomes. Core KOs were defined as orthologous groups present in all analyzed genomes, whereas accessory KOs were defined as those detected in only a subset of genomes. The distribution of shared and genome-specific KOs was visualized using UpSet plots (Lex et al. 2014).

For functional profiling, core KO identifiers were assigned to their corresponding KEGG level 2 functional categories, and the number of core KOs belonging to each functional category was determined for each genome. The relative abundance of each functional category was calculated by dividing the number of core KOs assigned to that category by the total number of core KOs. The 15 most abundant KEGG level 2 functional categories among the core KOs were identified based on their mean relative abundance across the four genomes and visualized using bar plots.

To characterize functional traits potentially associated with algal–bacterial interactions, KOs belonging to relevant KEGG pathways, including quorum sensing, biofilm formation, bacterial secretion systems, bacterial chemotaxis, flagellar assembly, and ABC transporters, were extracted from the annotation results. The presence and distribution of genes within these interaction-related pathways were compared across the four *Pseudomonas* genomes and visualized as a heatmap.

### 2.7 | Acridine Orange Staining and Fluorescence Microscopy

*C*. *reinhardtii* (4 × 10^6^ cells mL^−1^) and bacteria (1.6 × 10^9^ CFU mL^−1^) cells were mixed at an algal-to-bacterial cell ratio of ∼1:400. 200 µL of the mixed suspension was applied to glass slides coated with 0.5% (w/v) poly-L-lysine and incubated at room temperature for 10 min. The slides were then immersed in ice-cold methanol for 20 s for fixation and air-dried, following the protocol described by Aiyar et al. (2017). Subsequently, the slides were stained with 100 µg mL^−1^ acridine orange in 1% (v/v) acetic acid for 2 min, followed by gentle washing with deionized water. Fluorescence imaging was performed using an Axiophot fluorescence microscope (Carl Zeiss, Germany) equipped with filters (excitation: 450–490 nm; emission: 520 nm). Images were captured at 1000× magnification using an Axiocam ICc1 camera and Zen Blue software (Carl Zeiss).

### 2.8 | antiSMASH Analysis

The genomes of the three antagonistic *Pseudomonas* isolates and the reference *P. protegens* Pf-5 were analyzed using antiSMASH bacterial version 8.0.4 (Blin et al. 2025) to identify biosynthetic gene clusters (BGCs) involved in the production of secondary metabolites with the strictness set to “relaxed”.

### 2.9 | Statistical Analyses

Data were statistically analyzed using Microsoft Excel (Version 2408). Differences between treatments were assessed using a two-sided *t*-test assuming unequal variance (Welch’s *t*-test). Data are presented as means of standard deviation (SD) from three independent experiments.

## 3 | Results and Discussion

### 3.1 | Three *Pseudomonas* Isolates from the Rice Rhizosphere Antagonize ***Chlamydomonas***

Albeit many root associated rice bacteria have been studied (Edwards et al. 2015; Dastogeer et al. 2022), species identification is mostly missing. However, species identification is essential as the members of the same genus may have versatile metabolic capacity and exert distinct functions (Spiers et al. 2000; Nur Mawaddah et al. 2023; Sultana et al. 2024). In this study, bacterial strains were isolated from the rice rhizosphere (indicated by r) and rice soil (indicated by s) of an organic rice field in Italy (Figure 1a-c; Methods). A total of 12 bacterial strains were randomly selected due to their macroscopic characteristics (Table S1, Figure S2). All isolates exhibited circular colonies, most of which are opaque, whereas R-Bac-11s formed translucent colonies (Table S1, Figure S2). Five isolates derived from the rhizosphere, displaying colony colors from white to beige (Table S1, Figure S2). The remaining seven isolates from soil showed bright yellow, light yellow or white pigmentation (Table S1, Figure S2). Their molecular identification based on *16S rRNA* gene phylogeny revealed that the 12 strains belong to four genera, *Pseudomonas*, *Enterobacter,* the closely associated *Pantoea*, and *Paenarthrobacter* (Figures S3, S4; Table S2). All four genera have been reported to exhibit land plant growth-promoting properties (Shariati et al. 2017; Ngalimat et al. 2021; Salimi et al. 2023). Among them, the genus *Pseudomonas* was predominant, including eight isolates (Figure S3).

**FIGURE 1.**
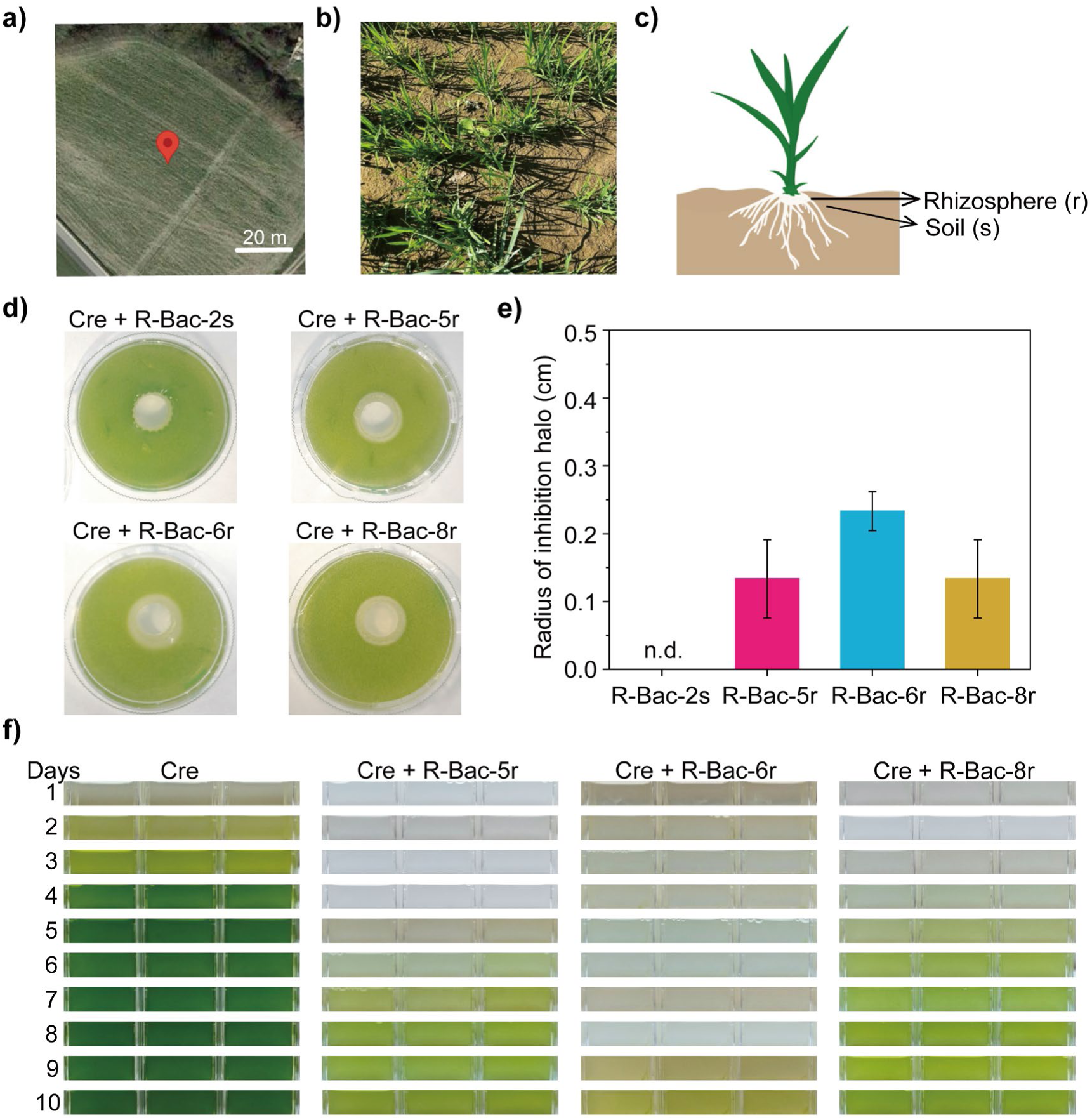
Three bacterial isolates from the rice rhizosphere exert antagonistic effects on *Chlamydomonas reinhardtii* (Cre). (a) Map showing the location of the rice field sampling site in Ottobiano, Italy. (b) Representative photograph of the rice cultivation field during the sampling period. (c) Schematic of a rice plant showing the rhizosphere (r) and surrounding soil (s), indicating the sources of bacterial isolation. (d) Growth inhibition agar plate assays of Cre with bacterial isolates from the rice field. Bacteria were inoculated on a circular area in the middle of the agar plate that was fully covered by an algal top layer (Methods). Images were taken after five days of growth. Bacteria R-Bac-5r, R-Bac-6r, and R-Bac-8r exhibited inhibitory effects on Cre growth. For further comparison, isolate R-Bac-2s, which showed no inhibition zone (commensal bacterium), is also presented. Two additional independent biological replicates are presented in Figure S5. (e) The radii of inhibition zones produced by R-Bac-2s and bacterial isolates R-Bac-5r, R-Bac-6r, and R-Bac-8r were measured after 5 days of incubation. The data are presented as mean ± standard deviation (n = 3). n.d., non-detectable. (f) Algal monocultures and liquid cocultures of Cre with the antagonistic bacteria R-Bac-5r, R-Bac-6r, or R-Bac-8r, photographed daily over 10 days. The organisms were cocultivated in TAP medium at a ratio of 1:250 (algae:bacteria). Two further independent biological replicates are shown in Figure S7a.

To determine if the 12 bacterial soil and rhizosphere isolates possess algicidal or growth promoting activities, they were screened against *C. reinhardtii* utilizing a standardized bipartite plate assay (Aiyar et al. 2017). Three bacteria (R-Bac-5r, R-Bac-6r, and R-Bac-8r) from the rhizosphere affected the growth of *C. reinhardtii* negatively (Figures 1d, S5). The other bacteria behaved commensally; they did not alter algal growth. The latter include R-Bac-2s (Figures 1d, S5) as well as R-Bac 1s, 3s, 4r, 7s, 9r, 10s, 11s and 12 s (Figure S6).

Based on the inhibition zones on agar plates, R-Bac-6r exerted the strongest growth inhibition (average of 0.23 ± 0.06 cm), followed by R-Bac-5r and R-Bac-8r that showed similar growth inhibition (averages of 0.13 ± 0.06 cm) (Figure 1e). All three newly identified antagonistic rhizosphere bacteria clustered within the *Pseudomonas* clade (Figure S3). Studies revealed diverse interactions between *Pseudomonas* bacteria and green algae. Under photoautotrophic conditions, cocultivation with symbiotic *Pseudomonas* sp. enhances the growth of *Chlorella vulgaris* (Guo and Tong 2014). *Pseudomonas gessardii* promotes the salt stress tolerance of *Chlorella sorokiniana* (Wang et al. 2024). By contrast, *Pseudomonas* sp. isolated from contaminated algal cultures slightly inhibits the growth of *C*. *reinhardtii* strain cc849 while increasing H_2_ accumulation (Li et al. 2013). *P. protegens* Pf-5, however, is a strong antagonist of *C. reinhardtii*, grown mixotrophically in the presence of acetate (Aiyar et al., 2017; Hotter et al. 2021; Rose et al. 2021).

The inhibitory effects of the three rice rhizosphere *Pseudomonas* isolates were further confirmed in liquid algal cocultures grown mixotrophically in Tris-Acetate-Phosphate (TAP) medium (Harris 1989) (Figures 1f, S7a). Algal cultures exhibited pronounced bleaching in the presence of the bacteria compared to algal monoculture controls, reflecting a stark lack of photosynthetic pigments (Figures 1f, S7a). The duration and severity of this bleaching mirrored the antagonism hierarchy observed on agar plates; cocultures exposed to R-Bac-6r remained severely bleached for a more extended period before any recovery, compared to the others.

In the next step, we analyzed the growth dynamics of all three bacterial strains and *C. reinhardtii* in TAP medium in mono- and cocultures (Figure 2). All used microorganisms can grow in the acetate-bearing TAP medium. While the cell densities of R-Bac-5r, and R-Bac-8r were not significantly different in mono- and cocultures, R-Bac-6r reached at some days a small but significant increase in the algal coculture compared to its monoculture (Figure 2a-c). This indicates that R-Bac-6r profits from the algae. In all cases the bacteria reached their highest levels on Day 2 in mono-and cocultures, before they declined (Figure 2a-c). As there were no significant differences in bacterial growth of R-Bac-5r, and R-Bac-8r in their mono- and cocultures together with the algae, these antagonistic bacteria seem not to profit from the algae. Such a behavior is known as amensialism (Faust and Raes 2012). One partner is negatively affected, while the other partner shows no effect.

**FIGURE 2.**
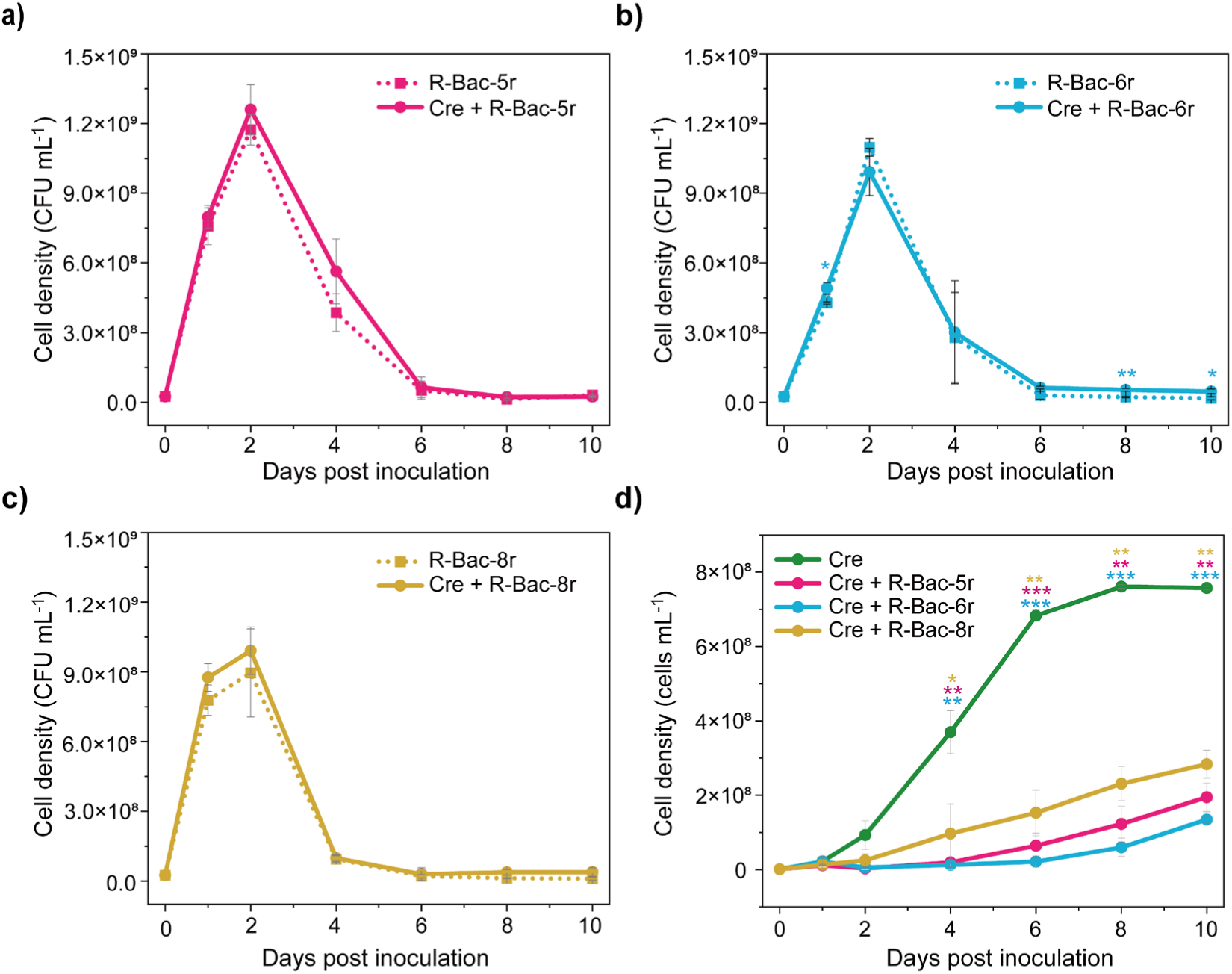
Growth dynamics of isolated antagonistic bacteria and *C*. *reinhardtii* (Cre) in mono- and cocultures. (a-c) Bacterial cell densities (Methods) of R-Bac-5r (a), R-Bac-6r (b), and R-Bac-8r (c), in mono- and coculture [see Figure 1f] with Cre over a 10-day period. (d) Algal cell densities (Methods) in mono-and coculture with R-Bac-5r, R-Bac-6r, and R-Bac-8r over a 10-day period. (a-d) Asterisks indicate statistically significant differences between monoculture and coculture as calculated by a two-sided *t*-test assuming unequal variance (Welch’s *t*-test). Data are presented as mean ± standard deviation. * *P*< 0.05, ** *P*< 0.01, *** *P*< 0.001. The absence of asterisk indicates no statistically significant difference.

To verify this further, we examined the cell number of *C. reinhardtii* in mono- and cocultures. The cell density of *C. reinhardtii* in coculture with R-Bac-5r, R-Bac-6r and R-Bac-8r decreased strongly in all cases, compared to an algal monoculture (Figure 2d). R-Bac-6r caused the strongest algal growth reduction followed by R-Bac-5r and R-Bac-8r. These results corroborate that all three *Pseudomonas* isolates exert antagonistic effects on *C*. *reinhardtii*.

We also examined the algal cell morphology in mono- and cocultures. In the presence of each of the three *Pseudomonas* bacteria, the algal oval cell shape changed. Notably, all *Pseudomonas* spp. investigated disrupted algal cell morphology, inducing cells to appear circular with a granular internal structure (Figures 3a, S7b).

**FIGURE 3.**
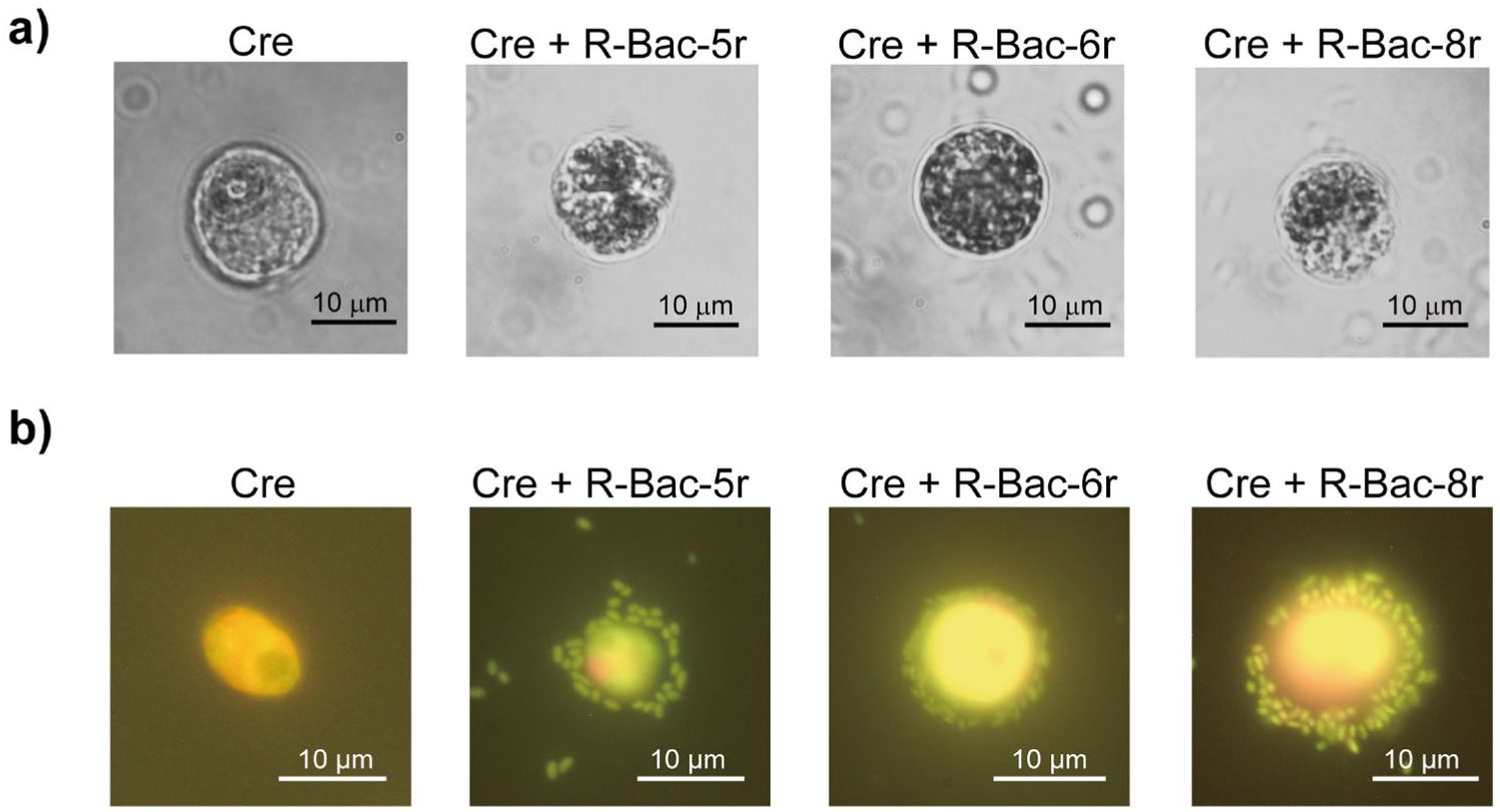
Antagonistic *Pseudomonas* rhizosphere bacteria change algal morphology and encircle algal cells. (a) Bright-field microscopy images of *C*. *reinhardtii* (Cre) cells in monoculture or cocultivated with R-Bac-5r, R-Bac-6r, and R-Bac-8r for 24 h. Two further independent biological replicates are shown in Figure S7b. (b) Acridine orange staining (Methods) visualizing the physical attachment of Cre and R-Bac-5r, R-Bac-6r, as well as R-Bac-8r after 10 min of incubation. Two further independent biological replicates are shown in Figure S7c.

To investigate potential physical interactions between *C*. *reinhardtii* and the antagonistic *Pseudomonas* spp., the cocultured microorganisms were stained with acridine orange and examined by fluorescence microscopy (Methods). All three antagonistic *Pseudomonas* spp. completely encircled the algal cells (Figures 3b, S7c). This close physical association may represent a prerequisite for transferring and injecting bacterial toxins efficiently into the algal cells, and to concentrate bacterial toxins. Thus, it was shown in case of *P. protegens* Pf-5 that the bacteria encircle the algal cells; they achieve a high local orfamide A concentration by this way needed to deflagellate the algal cells (Aiyar et al. 2017).

### 3.2 | Genomics Corroborates Antagonistic Effect of the *Pseudomonas* spp. On *Chlamydomonas* by Chemical Warfare

We performed whole-genome sequencing and annotation of the three antagonistic *Pseudomonas* spp. to determine their taxonomic identities at the species level. An overview of the genomic features of the three *Pseudomonas* spp. is presented in Table 1. The *P. protegens* Pf-5 genome was included for comparison. The respective genome sizes were 5,801,325 bp (R-Bac-5r), 6,562,028 bp (R-Bac-6r), and 5,672,562 bp (R-Bac-8r) and thus slightly smaller than *P. protegens* Pf-5 (7,074, 893 bp). All of them exhibit a high GC content ranging from 62% (*P. alloputida*) to 67% (*P. tohonis*). The whole-genome alignment-based phylogenetic relationships of the four *Pseudomonas* spp. with other related species obtained from the TYGS database were analyzed (Methods). Phylogenomic analysis classified the three rice-rhizosphere *Pseudomonas* spp. into three distinct clades (Figure 4). R-Bac-5r and R-Bac-8r belong to the same major clade, clustering with *P*. *alloputida* NMI24_14 and *P*. *asiatica* JCM 32716^T^, respectively, whereas R-Bac-6r clustered in a separate clade with *P*. *tohonis* TUM18999^T^ (Figure 4). Minimal standards for the use of genome data for the taxonomy of prokaryotes, based on the combination of *16S rRNA* gene similarity and overall genome related indexes (average nucleotide identity, ANI, and digital DNA-DNA hybridization, dDDH) have been proposed (Chun et al. 2018). The species delineation thresholds are 95– 96% for ANI (Richter and Rosselló-Móra 2009), 70% for dDDH (Goris et al. 2007), and 98.7% for *16S rRNA* gene similarity (Kim et al. 2014). The ANI, dDDH, and *16S rRNA* gene similarity values between each investigated strain and their closest reference strains being R-Bac-5r and *P*. *alloputida* NMI24_14, R-Bac-6r and *P*. *tohonis* TUM18999^T^, as well as R-Bac-8r and *P*. *asiatica* JCM 32716^T^, were all higher than 96% (ANI), 75% (dDDH), and 99% (*16S rRNA*, Figure 5). Thus, they exceed the established species delineation thresholds. Therefore, our results identify R-Bac-5r, R-Bac-6r, and R-Bac-8r as *P*. *alloputida*, *P*. *tohonis*, and *P*. *asiatica*, respectively.

**Table 1.** General genomic features of the isolated *Pseudomonas* spp. and *P. protegens* Pf-5.

| **Bacteria** | ***P. alloputida***<br>**R-Bac-5r** | ***P. tohonis***<br>**R-Bac-6r** | ***P. asiatica***<br>**R-Bac-8r** | ***P. protegens***<br>**Pf-5<sup>a</sup>** |
|---|---|---|---|---|
| Genome size (bp) | 5,801,325 | 6,562,028 | 5,672,562 | 7,074,893 |
| GC content (%) | 62 | 67 | 63 | 63 |
| Total contigs | 152 | 106 | 139 | 1 |
| Largest contig (bp) | 449,201 | 699,992 | 428,151 | 7,074,893 |
| Contig N50 (bp)<sup>b</sup> | 227,023 | 300,615 | 269,028 | 7,074,893 |
| Coding sequences (CDSs) | 5,263 | 5,919 | 5,172 | 6,276 |
| Contamination (%)<sup>c</sup> | 0.19 | 0.0 | 0.2 | 0.48 |
| Completeness (%) | 100 | 100 | 100 | 100 |

<sup>a</sup>The *P*. *protegens* Pf-5 genome (GenBank assembly accession GCA\_000012265.1) was ob-tained from the NCBI Assembly database.
<sup>b</sup>Contig N50 is the length of the shortest contig in the set of longest contigs that collectively represent at least 50% of the total assembly size.
<sup>c</sup>Contamination is calculated by CheckM2, which estimates the proportion of non-target or erroneously duplicated genomic content in an assembly using a gradient boosting–based prediction model.

**FIGURE 4.**
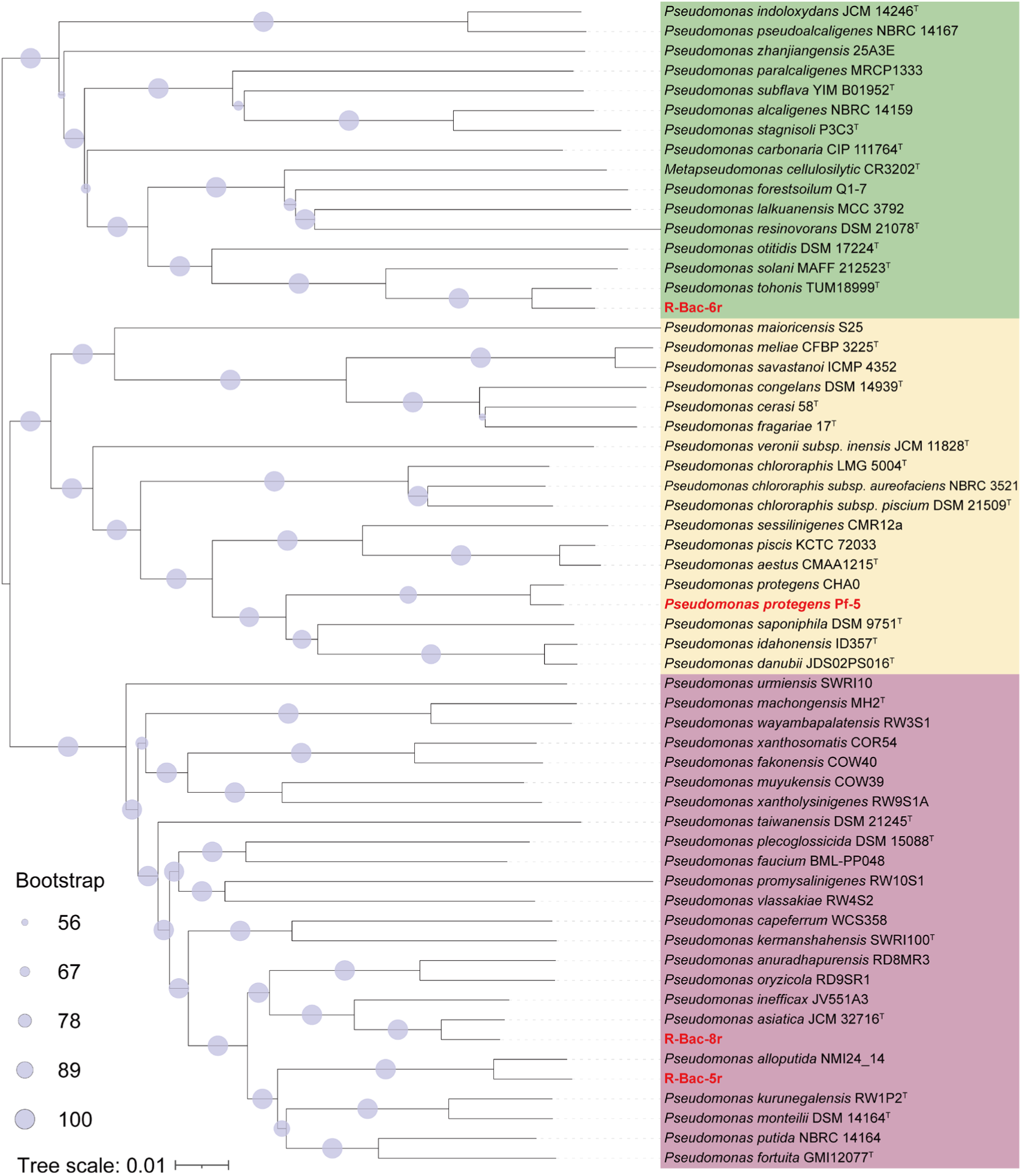
Whole-genome phylogenetic analysis of the antagonistic rice rhizosphere isolates R-Bac-5r, R-Bac-6r, and R-Bac-8r compared to the antagonist *Pseudomonas protegens* Pf-5 (all indicated in red colour). Phylogenetic relationships were inferred using the Genome BLAST Distance Phylogeny (GBDP) method implemented in the Type (Strain) Genome Server (TYGS). The phylogenetic tree was reconstructed with FastME v2.1.6.1 from GBDP-derived intergenomic distances, with branch lengths representing the GBDP distance formula d5 (Methods).

**FIGURE 5.**
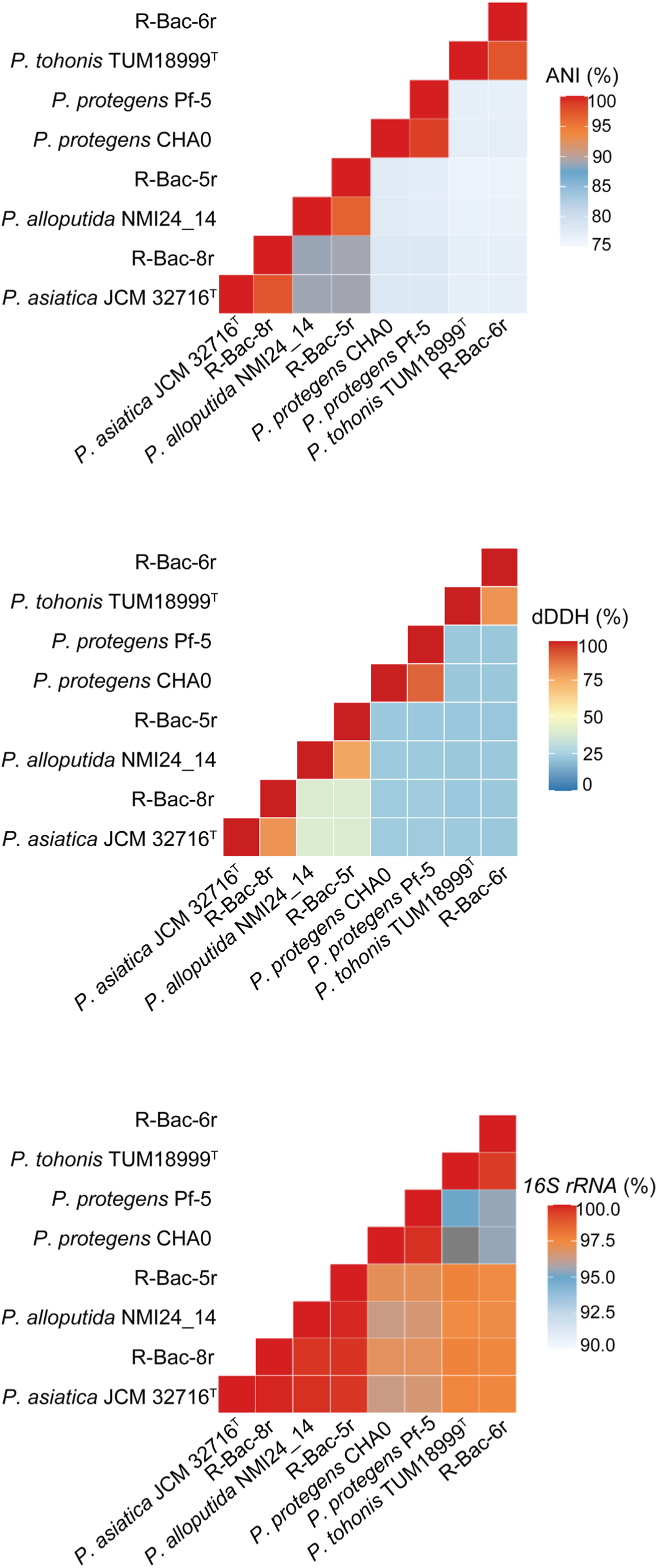
Species delineation of three antagonistic rice rhizosphere isolates, R-Bac-5r, R-Bac-6r, and R-Bac-8r, within the genus *Pseudomonas* identify them as *P. alloputida*, *P. tohonis* and *P. asiatica*, respectively. The heatmap displays pairwise comparisons of average nucleotide identity (ANI), digital DNA–DNA hybridization (dDDH), and *16S rRNA* gene sequence identity among the three isolates and five additional *Pseudomonas* genomes (Methods). Species boundaries were assessed using established thresholds of 95–96% for ANI (Richter and Rosselló-Móra 2009), 70% for dDDH (Goris et al. 2007), and 98.7% for 16S rRNA gene sequence similarity (Kim et al. 2014).

*Pseudomonas* is one of the most diverse and ubiquitous bacterial genera (Peix et al. 2018). *P*. *alloputida*, *P*. *tohonis*, and *P. asiatica* were classified as new species in recent years (Keshavarz-Tohid et al. 2019; Tohya et al. 2019; Yamada et al. 2021). Currently, studies on their interactions with microalgae remain lacking. The identification of these three *Pseudomonas* spp. suggests that antagonistic activity toward *C*. *reinhardtii* is not limited to *P*. *protegens* Pf-5 but is distributed across multiple *Pseudomonas* species. *P. protegens* and *P. tohonis* have been characterized, for example, as biocontrol agents against phytopathogenic bacteria (Wang et al. 2023). *P*. *asiatica* could act as an efficient phosphate-solubilizing bacterium (Wang et al. 2022). In general, *Pseudomonas* spp. are well known to act as a biocontrol agents that are beneficial for growth of crops, including rice (Mercado-Blanco and Bakker 2007). Interestingly, aqueous extracts of *Chlamydomonas* sp. stimulate rice plant development as well (Lamb et al., 2023). It seems that the antagonistic rhizosphere *Pseudomonas* spp. suppress and possibly lyse microalgae, thereby increasing nutrient availability for the host rice plant. As the *C. reinhardtii* phycosphere and *Arabidopsis thaliana* roots recruit similar bacterial communities (Durán et al. 2022), the microalga may represent a direct competitor in the rice rhizosphere, triggering chemical suppression by the native root bacteria.

To better understand the molecular mechanism how the rhizosphere *Pseudomonas* spp. may exert their algal antagonism, the KEGG database was used to analyze their functions. Nearly 2,000 KEGG Orthology (KO) groups were shared across the three antagonistic *Pseudomonas* spp., with *P*. *protegens* Pf-5 included for comparison (Figure S8a). Core KO categories were determined highlighting the conservation of fundamental biological functions among the four *Pseudomonas* genomes (Figure S8b). Specifically, pathways associated with algal–bacterial interactions were depicted among the four *Pseudomonas* spp. Remarkably, these species encode gene repertoires associated with pathways involved in microbial interactions, embracing chemical warfare (Figure 6a). These include ABC (ATP-binding cassette) transporters by which bacteria may expel their toxins (Beis and Rebuffat 2019). ABC transporters are closely related to the development of biofilm formation in *Pseudomonas* spp. whose genes were also found and contribute to cell-to-cell interactions (García-Bayona and Comstock 2018). Bacterial flagella-driven swimming motility and chemotaxis are considered key determinants for locating algal cells and initiating surface attachment (Forget et al. 2026), thereby promoting close physical adhesion. This interpretation is consistent with the microscopy observations showing that all four *Pseudomonas* strains closely surrounded *C*. *reinhardtii* cells (Figure 3b). Collectively, these findings suggest that the four *Pseudomonas* spp. possess conserved genetic capacities to support algal antagonism.

**FIGURE 6.**
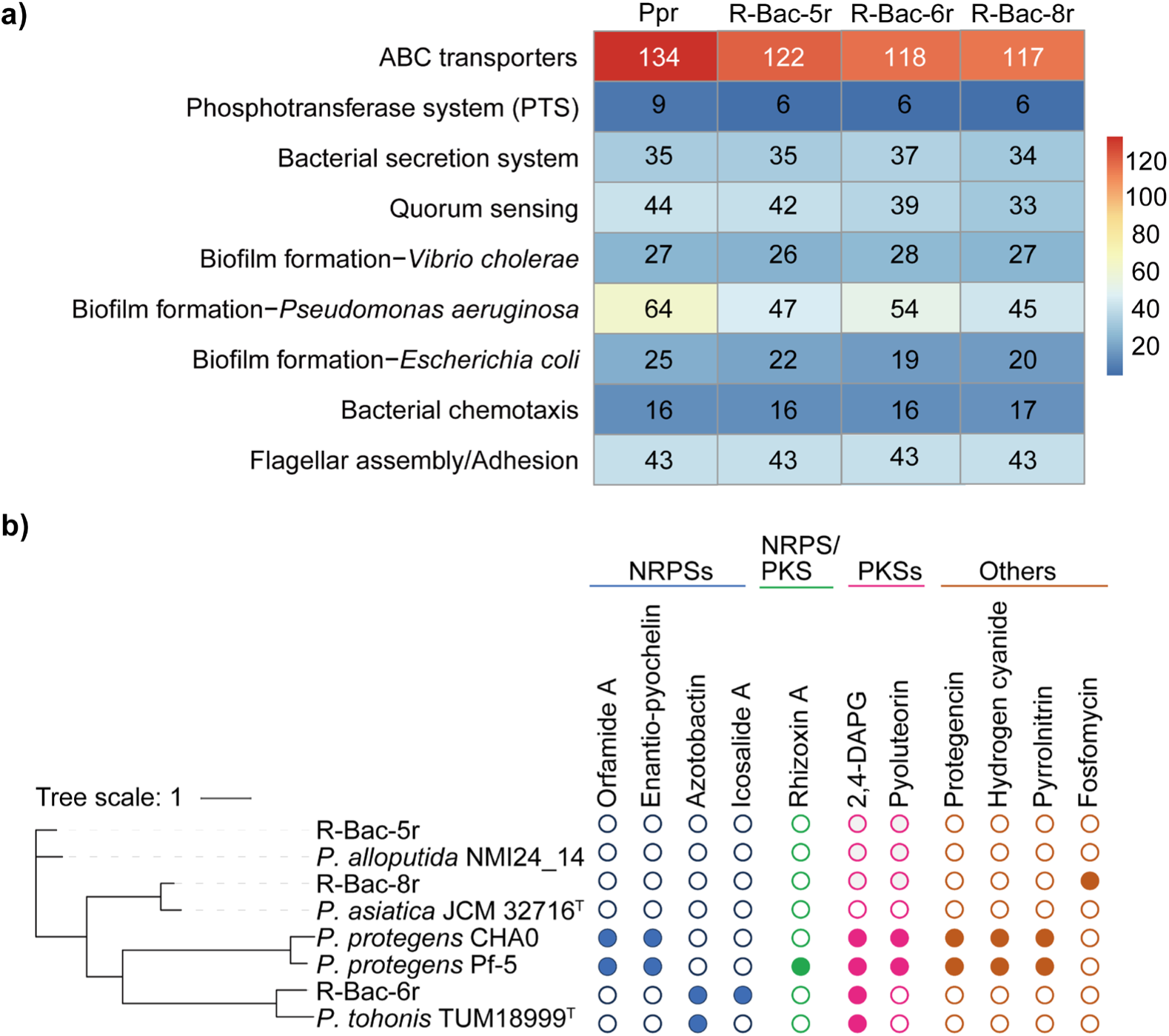
Genomic comparison of antagonistic *Pseudomonas* spp. revealing chemical warfare strategies (a) Heatmap, illustrating the presence of genes associated with selected metabolic path-ways potentially involved in algal-bacterial interactions in *P*. *protegens* (Ppr), R-Bac-5r, R-Bac-6r, as well as R-Bac-8r. (b) Comparison of phylogeny and potential secondary metabolites of biosynthetic gene clusters (filled circles = presence of a gene or gene cluster). Predicted secondary metabolites represent potential bioactive compounds that may contribute to bacterial antagonism and inhibition of *C*. *reinhardtii* growth. NRPS, non-ribosomal peptide synthetase; PKS, polyketide synthase. Orfamide A, rhizoxin A, 2,4-DAPG, pyoluteorin, protegencin and pyrrolnitrin are already known for their toxic effects on *C. reinhardtii* (Aiyar et al. 2017; Hotter et al. 2021; Rose et al. 2021).

Differences in the synthesis of toxic secondary metabolites may cause the variation in antagonistic activity among the isolates. To identify potential secondary metabolites, the genomes were mined for BGCs associated with potential antialgal compounds using antiSMASH. The analysis of *P. protegens* Pf-5 and its closest type strain was used for comparison and showed the presence of known secondary metabolites, including toxic ones (Aiyar et al. 2017; Hotter et al. 2021; Rose et al. 2021). Especially interesting is the predicted presence of the phenolic secondary metabolite DAPG in the strongest antagonist, R-Bac-6r and its closest type strain (Figure 6b). DAPG, a polyketide produced by some *Pseudomonas* strains, exhibits a broad-spectrum inhibitory activity against bacteria. It has become, for example, a microbial bio-product in agrochemical applications against tomato bacterial wilt (Suresh et al. 2022). Notably, rather than acting as a plant protectant, DAPG exerts a toxic influence on *C. reinhardtii*, inhibiting its growth and altering its cell morphology (Rose et al. 2021). Therefore, the presence of 2,4-DAPG BGC in R-Bac-6r (*P. tohonis*) suggests that this metabolite, either alone or in combination with additional uncharacterized toxins, contributes to the antagonistic activity. A non-ribosomal peptide synthetase (NRPS)-type BGC responsible for icosalide A biosynthesis was also detected in R-Bac-6r (Figure 6b). Icosalides have antimicrobial activity against some bacteria (Boros et al. 2006) but are not known so far to potentially antagonize algae. The genome of R-Bac-8r contains the BGC predicted to direct fosfomycin biosynthesis, which might interfere with algal plastid division (Matsumoto et al. 2012); it is absent in its closest type strain. No known BGCs were identified in the genomes of R-Bac-5r or its closest type strain (Figure 6b). Despite this absence of established BGCs, R-Bac-5r exhibits robust antagonistic activity against *C. reinhardtii*, suggesting that this strain relies on uncharacterized or novel secondary metabolites to mediate algal suppression. Finally, the azotobactin biosynthetic gene cluster was also identified in both R-Bac-6r and its closest type strain genomes (Figure 6b). Azotobactin conventionally functions as a siderophore and can supply nitrogen to support the growth of green algae (Villa et al. 2014). In this scenario, the rice plant may profit instead.

## 4 | Conclusion

In summary, our results identify three rhizosphere-derived *Pseudomonas* isolates with antagonistic activity against *C*. *reinhardtii*. Comparative genomic analyses suggest that common pathways for microbial interactions and different - partially unknown - secondary metabolites, including the presence of the DAPG biosynthetic gene cluster in *P*. *tohonis* R-Bac-6r, contribute to strain-specific antagonistic activity. Interestingly, *Pseudomonas* spp. including the toxin DAPG are known to protect crop plants while they are antagonistic against the algae. These findings expand our understanding of the genetic diversity underlying *Pseudomonas*–microalgal antagonistic interactions and highlight rice rhizosphere-associated bacteria as an important reservoir of functional traits that influence entire soil population dynamics.

## Acknowledgements

We thank Trang Vuong for valuable discussions and for proof-reading the manuscript. MM received funding from the DFG within the Germany’s Excellence Strategy – EXC 2051 – Project ID 390713860. XY is supported with a postdoctoral scholarship from the Chinese Academy of Sciences (CAS) Scholarship.

## Data availability statement

The data that support the findings of this study are available in this manuscript and its Supporting Information.

The Illumina short-read sequencing data for the *Pseudomonas* isolates R-Bac-5r, R-Bac-6r, and R-Bac-8r have been deposited under the accession number PRJNA1507846. Reviewer link: https://dataview.ncbi.nlm.nih.gov/object/PRJNA1507846?reviewer=hmda234sdoppiph404hd k5spl

## Ethics statement

All experimental work complied with prevailing local, national and international regulations and conventions, and normal scientific ethical practices.

## Author contributions

PS and MM designed the project. XY, PS, BV and IUH performed experiments, and all authors contributed to the analysis and interpretation of results. XY and MM wrote the paper with inputs from all coauthors.

## Supporting Information

Additional supporting information can be found online in the Supporting Information section.

**Table S1:** Overview of 12 bacterial isolates from either the rhizosphere (r) or the surrounding soil (s) from a rice field in Ottobiano, Italy.

**Table S2:** *16S rRNA* gene sequence dataset obtained from public databases.

**Figure S1:** Calibration of optical density measurements for three antagonistic bacterial isolates.

**Figure S2:** Colony morphology of 12 bacterial strains isolated from rice soil (s) or the rice rhizosphere (r).

**Figure S3:** Phylogenetic relationships of bacterial isolates from the rice field based on *16S rRNA* gene sequences (Methods).

**Figure S4:** Amplified *16S rRNA* gene sequences (Methods) of 12 bacterial isolates from the rice field soil (s) or the rice rhizosphere (r), presented in the 5′–3′ direction.

**Figure S5:** Antagonistic rice rhizosphere isolates R-Bac-5r, R-Bac-6r, and R-Bac-8r compared to the commensal R-Bac-2s.

**Figure S6:** Screening of rice field bacterial isolates against *C*. *reinhardtii* (Cre).

**Figure S7:** Cocultivation, cellular morphology and physical association of *C*. *reinhardtii* (Cre) with antagonistic *Pseudomonas* spp.

**Figure S8:** Comparative genomic analysis of *Pseudomonas protegens* (Ppr) and antagonistic rice rhizosphere isolates R-Bac-5r, R-Bac-6r, and R-Bac-8r.

## REFERENCES

1. Abate, R., Oon, Y. L., Oon, Y. S., et al. 2024. “Diverse interactions between bacteria and microalgae: A review for enhancing harmful algal bloom mitigation and biomass processing efficiency.” Heliyon 10(17):e36503. 10.1016/j.heliyon.2024.e36503.

2. Aiyar, P., Schaeme, D., García-Altares, M., et al. 2017. “Antagonistic bacteria disrupt calcium homeostasis and immobilize algal cells.” Nature Communications 8:1756. 10.1038/s41467-017-01547-8.

3. Bais, H. P., Weir, T. L., Perry, L. G., Gilroy, S., and Vivanco, J. M. 2006. “The role of root exudates in rhizosphere interactions with plants and other organisms.” Annual Review of Plant Biology 57:233–266. 10.1146/annurev.arplant.57.032905.105159.

4. Barrett, M. R., and Koch, A. R. 1982. “Effects of ammonium sulfate and urea on the growth of Chlorophycean algae from rice fields.” Journal of Environmental Quality 11:187–191. 10.2134/jeq1982.00472425001100020009x.

5. Beis, K., and Rebuffat, S. 2019. “Multifaceted ABC transporters associated to microcin and bacteriocin export.” Research in Microbiology 170:399–406. 10.1016/j.resmic.2019.07.002.

6. Blin, K., Shaw, S., Vader, L., et al. 2025. “antiSMASH 8.0: extended gene cluster detection capabilities and analyses of chemistry, enzymology, and regulation.” Nucleic Acids Research 53:W32–W38. 10.1093/nar/gkaf334.

7. Boros, C., Smith, C. J., Vasina, Y., et al. 2006. “Isolation and identification of the icosalides--cyclic peptolides with selective antibiotic and cytotoxic activities.” The Journal of Antibiotics 59:486–494. 10.1038/ja.2006.68.

8. Breidenbach, B., Pump, J., and Dumont, M. G. 2016. “Microbial community structure in the rhizosphere of rice plants.” Frontiers in Microbiology 6:1537. 10.3389/fmicb.2015.01537.

9. Bunbury, F., Deery, E., Sayer, A. P. et al. 2022. “Exploring the onset of B_12_-based mutualisms using a recently evolved *Chlamydomonas* auxotroph and B_12_-producing bacteria.” Environmental Microbiology 24:3134–3147. 10.1111/1462-2920.16035.

10. Burgunter-Delamare, B., Shetty, P., Vuong, T., and Mittag, M. 2024. “Exchange or eliminate: The secrets of algal-bacterial relationships.” Plants 13:829. 10.3390/plants13060829.

11. Calatrava, V., Hom, E. F. Y., Guan, Q., Llamas, A., Fernández, E., and Galván, A. 2024. “Genetic evidence for algal auxin production in *Chlamydomonas* and its role in algal-bacterial mutualism.” iScience 27:108762. 10.1016/j.isci.2023.108762.

12. Calatrava, V., Tejada-Jimenez, M., Sanz-Luque, E., Fernandez, E., Galvan, A., and Llamas, A. 2023. “*Chlamydomonas reinhardtii*, a reference organism to study algal-microbial interactions: why can’t they be friends?” Plants 12:788. 10.3390/plants12040788.

13. Carrasco Flores, D., Hotter, V., Vuong, T., et al. 2024. “A mutualistic bacterium rescues a green alga from an antagonist.” Proceedings of the National Academy of Sciences of the United States of America 121: e2401632121. 10.1073/pnas.2401632121.

14. Chaumeil, P. A., Mussig, A. J., Hugenholtz, P., and Parks, D. H. 2019. “GTDB-Tk: a toolkit to classify genomes with the Genome Taxonomy Database.” Bioinformatics 36:1925–1927. 10.1093/bioinformatics/btz848.

15. Chen, S., Zhou, Y., Chen, Y., and Gu, J. 2018. “fastp: an ultra-fast all-in-one FASTQ preprocessor.” Bioinformatics 34:884–890. 10.1093/bioinformatics/bty560.

16. Chklovski, A., Parks, D. H., Woodcroft, B. J., and Tyson, G. W. 2023. “CheckM2: a rapid, scalable and accurate tool for assessing microbial genome quality using machine learning.” Nature Methods 20:1203–1212. 10.1038/s41592-023-01940-w.

17. Chun, J., Oren, A., Ventosa, A., et al. 2018. “Proposed minimal standards for the use of genome data for the taxonomy of prokaryotes.” International Journal of Systematic and Evolutionary Microbiology 68:461–466. 10.1099/ijsem.0.002516.

18. Cirri, E., and Pohnert, G. 2019. “Algae-bacteria interactions that balance the planktonic microbiome.” New Phytologist 223:100–106. 10.1111/nph.15765.

19. Dastogeer, K. M. G., Yasuda, M., and Okazaki, S. 2022. “Microbiome and pathobiome analyses reveal changes in community structure by foliar pathogen infection in rice.” Frontiers in Microbiology 13:949152. 10.3389/fmicb.2022.949152.

20. Ding, L. J., Cui, H. L., Nie, S. A., Long, X. E., Duan, G. L., and Zhu, Y. G. 2019. “Microbiomes inhabiting rice roots and rhizosphere.” FEMS Microbiology Ecology 95:fiz040. 10.1093/femsec/fiz040.

21. Durán, P., Flores-Uribe, J., Wippel, K., et al. 2022. “Shared features and reciprocal complementation of the *Chlamydomonas* and *Arabidopsis* microbiota.” Nature Communications 13:406. 10.1038/s41467-022-28055-8.

22. Edwards, J., Johnson, C., Santos-Medellín, C., et al. 2015. “Structure, variation, and assembly of the root-associated microbiomes of rice.” Proceedings of the National Academy of Sciences of the United States of America 112:E911–E920. 10.1073/pnas.1414592112.

23. Faust, K., and Raes, J. 2012. “Microbial interactions: From networks to models.” Nature Reviews Microbiology 10: 538–550. 10.1038/nrmicro2832.

24. Felsenstein, J. 1981. “Evolutionary trees from DNA sequences: a maximum likelihood approach.” Journal of Molecular Evolution 17:368–376. 10.1007/BF01734359.

25. Field, C. B., Behrenfeld, M. J., Randerson, J. T., and Falkowski, P. 1998. “Primary production of the biosphere: integrating terrestrial and oceanic components.” Science 281:237–240. 10.1126/science.281.5374.237.

26. Forget, M., Müller, O., and Stocker, R. 2026. “The ecology of bacterial attachment to phytoplankton.” Nature Microbiology 11:1180–1192. 10.1038/s41564-026-02287-6.

27. García-Bayona, L., and Comstock, L. E. 2018. “Bacterial antagonism in host-associated microbial communities.” Science 361:eaat2456. 10.1126/science.aat2456.

28. Ghasemi, Y., Rasoul-Amini, S., Morowvat, M. H., et al. 2008. “Characterization of hydrocortisone biometabolites and *18S rRNA* gene in *Chlamydomonas reinhardtii* cultures.” Molecules 13:2416–2425. 10.3390/molecules13102416.

29. Goris, J., Konstantinidis, K. T., Klappenbach, J. A., Coenye, T., Vandamme, P., and Tiedje, J. M. 2007. “DNA–DNA hybridization values and their relationship to whole-genome sequence similarities.” International Journal of Systematic and Evolutionary Microbiology 57:81–91. 10.1099/ijs.0.64483-0.

30. Guo, Z., and Tong, Y. W. 2014. “The interactions between *Chlorella vulgaris* and algal symbiotic bacteria under photoautotrophic and photoheterotrophic conditions.” Journal of Applied Phycology 26:1483–1492. 10.1007/s10811-013-0186-1.

31. Harris, E. H. 1989. The Chlamydomonas sourcebook. A comprehensive guide to biology and laboratory use. Academic Press.

32. Hori, T., Noll, M., Igarashi, Y., Friedrich, M. W., and Conrad, R. 2007. “Identification of acetate-assimilating microorganisms under methanogenic conditions in anoxic rice field soil by comparative stable isotope probing of RNA.” Applied and Environmental Microbiology 73:101–109. 10.1128/aem.01676-06.

33. Hotter, V., Zopf, D., Kim, H. J., et al. 2021. “A polyyne toxin produced by an antagonistic bacterium blinds and lyses a Chlamydomonad alga.” Proceedings of the National Academy of Sciences of the United States of America 118:e2107695118. 10.1073/pnas.2107695118.

34. Hussain, Q., Pan, G. X., Liu, Y. Z., et al. 2012. “A Microbial community dynamics and function associated with rhizosphere over periods of rice growth.” *Plant*, Soil and Environment 58:55–61. 10.17221/390/2010-PSE.

35. Islam, M. M., Jana, S. K., Sengupta, S., and Mandal, S. 2024. “Impact of rhizospheric microbiome on rice cultivation.” Current Microbiology 81:188. 10.1007/s00284-024-03703-y.

36. Keshavarz-Tohid, V., Vacheron, J., Dubost, A., et al. 2019. “Genomic, phylogenetic and catabolic re-assessment of the *Pseudomonas putida* clade supports the delineation of *Pseudomonas alloputida* sp. nov., *Pseudomonas inefficax* sp. nov., *Pseudomonas persica* sp. nov., and *Pseudomonas shirazica* sp. nov.” Systematic and Applied Microbiology 42:468–480. 10.1016/j.syapm.2019.04.004.

37. Kim, M., Oh, H. S., Park, S. C., and Chun, J. 2014. “Towards a taxonomic coherence between average nucleotide identity and *16S rRNA* gene sequence similarity for species demarcation of prokaryotes.” International Journal of Systematic and Evolutionary Microbiology 64:346–351. 10.1099/ijs.0.059774-0.

38. Kumar, S., Stecher, G., Suleski, M., Sanderford, M., Sharma, S., and Tamura, K. 2024. “MEGA12: Molecular evolutionary genetic analysis version 12 for adaptive and green computing.” Molecular Biology and Evolution 41:msae263. 10.1093/molbev/msae263.

39. Lamb, T. I., Berghahn, E., Pita, F. M., et al. 2023. “Isolation and selection of microalgae capable of stimulating rice plant development and seed production.” Algal Research 74:103203. 10.1016/j.algal.2023.103203.

40. Letunic, I., and Bork, P. 2024. “Interactive Tree of Life (iTOL) v6: recent updates to the phylogenetic tree display and annotation tool.” Nucleic Acids Research 52:W78–W82. 10.1093/nar/gkae268.

41. Lex, A., Gehlenborg, N., Strobelt, H., Vuillemot, R., and Pfister, H. 2014. “UpSet: visualization of intersecting sets.” IEEE Transactions on Visualization and Computer Graphics 20:1983–1992. 10.1109/TVCG.2014.2346248.

42. Li, X., Huang, S., Yu, J., Wang, Q., and Wu, S. 2013. “Improvement of hydrogen production of *Chlamydomonas reinhardtii* by co-cultivation with isolated bacteria.” International Journal of Hydrogen Energy 38:10779–10787. 10.1016/j.ijhydene.2013.02.102.

43. Lingappa, U. F., Borrego, S. D., Sindermann, E. S., et al. 2026. “Field strains of the unicellular alga *Chlamydomonas reinhardtii* exhibit multicellular characteristics that shape their interactions.” The ISME Journal 20:wrag202. 10.1093/ismejo/wrag202.

44. Liu, D., Zhang, Y., Fan, G., et al. 2022. “IPGA: A handy integrated prokaryotes genome and pan-genome analysis web service.” iMeta 1:e55. 10.1002/imt2.55.

45. Matsumoto, H., Takechi, K., Sato, H., Takio, S., and Takano, H. 2012. “Treatment with antibiotics that interfere with peptidoglycan biosynthesis inhibits chloroplast division in the desmid *Closterium*.” PLoS ONE 7:e40734. 10.1371/journal.pone.0040734.

46. Meier-Kolthoff, J. P., and Göker, M. 2019. “TYGS is an automated high-throughput platform for state-of-the-art genome-based taxonomy.” Nature Communications 10:2182. 10.1038/s41467-019-10210-3.

47. Mercado-Blanco, J., and Bakker, P. A. 2007. “Interactions between plants and beneficial *Pseudomonas* spp.: exploiting bacterial traits for crop protection.” Antonie van Leeuwenhoek 92:367–389. 10.1007/s10482-007-9167-1.

48. Moriya, Y., Itoh, M., Okuda, S., Yoshizawa, A. C., and Kanehisa, M. 2007. “KAAS: an automatic genome annotation and pathway reconstruction server.” Nucleic Acids Research 35:W182–W185. 10.1093/nar/gkm321.

49. Ngalimat, M. S., Mohd Hata, E., Zulperi, D., et al. 2021. “Plant growth-promoting bacteria as an emerging tool to manage bacterial rice pathogens.” Microorganisms 9:682. 10.3390/microorganisms9040682.

50. Nur Mawaddah, S., AW, M. Z., and Sapak, Z. 2023. “The potential of *Pseudomonas fluorescens* as biological control agent against sheath blight disease in rice: a systematic review.” Food Research 7:46–56. 10.26656/fr.2017.7(S2).11.

51. Overbeek, R., Olson, R., Pusch, G. D., et al. 2014. “The SEED and the Rapid Annotation of microbial genomes using Subsystems Technology (RAST).” Nucleic Acids Research 42:D206–D214. 10.1093/nar/gkt1226.

52. Peix, A., Ramírez-Bahena, M. H., and Velázquez, E. 2018. “The current status on the taxonomy of *Pseudomonas* revisited: An update.” Infection, Genetics and Evolution 57:106–116. 10.1016/j.meegid.2017.10.026.

53. Perrineau, M. M., Gross, J., Zelzion, E., et al. 2014. “Using natural selection to explore the adaptive potential of *Chlamydomonas reinhardtii*.” PLoS ONE 9:e92533. 10.1371/journal.pone.0092533.

54. Richter, M., and Rosselló-Móra, R. 2009. “Shifting the genomic gold standard for the prokaryotic species definition.” Proceedings of the National Academy of Sciences of the United States of America 106:19126–19131. 10.1073/pnas.0906412106.

55. Rose, M. M., Scheer, D., Hou, Y., et al. 2021. “The bacterium *Pseudomonas protegens* antagonizes the microalga *Chlamydomonas reinhardtii* using a blend of toxins.” Environmental Microbiology 23: 5525–5540. 10.1111/1462-2920.15700.

56. Salimi, F., Khorshidi, M., Amirahmadi, F., and Amirahmadi, A. 2023. “Effectiveness of phosphate and zinc solubilizing *Paenarthrobacter nitroguajacolicus* P1 as halotolerant rhizobacterium with growth-promoting activity on *Pistacia vera* L.” Current Microbiology 80:336. 10.1007/s00284-023-03448-0.

57. Shariati J, V., Malboobi, M. A., Tabrizi, Z., Tavakol, E., Owlia, P., and Safari, M. 2017. “Comprehensive genomic analysis of a plant growth-promoting rhizobacterium *Pantoea agglomerans* strain P5.” Scientific Reports 7:15610. 10.1038/s41598-017-15820-9.

58. Spiers, A. J., Buckling, A., and Rainey, P. B. 2000. “The causes of *Pseudomonas* diversity.” Microbiology 146:2345–2350. 10.1099/00221287-146-10-2345.

59. Sultana, R., Jashim, A. I. I., Islam, S. M. N., Rahman, M. H., and Haque, M. M. 2024. “Bacterial endophyte *Pseudomonas mosselii* PR5 improves growth, nutrient accumulation, and yield of rice (*Oryza sativa* L.) through various application methods.” BMC Plant Biology 24:1030. 10.1186/s12870-024-05649-6.

60. Suresh, P., Rekha, M., Gomathinayagam, S., et al. 2022. “Characterization and assessment of 2, 4-Diacetylphloroglucinol (DAPG)-producing *Pseudomonas fluorescens* VSMKU3054 for the management of tomato bacterial wilt caused by *Ralstonia solanacearum*.” Microorganisms 10:1508. 10.3390/microorganisms10081508.

61. Teplitski, M., Chen, H., Rajamani, S., et al. 2004. “*Chlamydomonas reinhardtii* secretes compounds that mimic bacterial signals and interfere with quorum sensing regulation in bacteria.” Plant Physiology 134:137–146. 10.1104/pp.103.029918.

62. Tohya, M., Watanabe, S., Teramoto, K., et al. 2019. “*Pseudomonas asiatica* sp. nov., isolated from hospitalized patients in Japan and Myanmar.” International Journal of Systematic and Evolutionary Microbiology 69:1361–1368. 10.1099/ijsem.0.003316.

63. Villa, J. A., Ray, E. E., and Barney, B. M. 2014. “*Azotobacter vinelandii* siderophore can provide nitrogen to support the culture of the green algae *Neochloris oleoabundans* and *Scenedesmus* sp. BA032.” FEMS Microbiology Letters 351:70–77. 10.1111/1574-6968.12347.

64. Wang, L., Zhou, F., Zhou, J., et al. 2022. “Genomic analysis of *Pseudomonas asiatica* JP233: An efficient phosphate-solubilizing bacterium.” Genes 13:2290. 10.3390/genes13122290.

65. Wang, S., Hu, M., Chen, H., et al. 2023. “*Pseudomonas forestsoilum* sp. nov. and *P*. *tohonis* biocontrol bacterial wilt by quenching 3-hydroxypalmitic acid methyl ester.” Frontiers in Plant Science 14:1193297. 10.3389/fpls.2023.1193297.

66. Wang, T., Li, D., Tian, X., et al. 2024. “Mitigating salinity stress through interactions between microalgae and different forms (free-living & alginate gel-encapsulated) of bacteria isolated from estuarine environments.” Science of The Total Environment 926:171909. 10.1016/j.scitotenv.2024.171909.

67. Wick, R. R., Judd, L. M., Gorrie, C. L., and Holt, K. E. 2017. “Unicycler: Resolving bacterial genome assemblies from short and long sequencing reads.” PLoS Computational Biology 13: e1005595. 10.1371/journal.pcbi.1005595.

68. Yamada, K., Sasaki, M., Aoki, et al. 2021. “*Pseudomonas tohonis* sp. nov., isolated from the skin of a patient with burn wounds in Japan.” International Journal of Systematic and Evolutionary Microbiology 71:005115. 10.1099/ijsem.0.005115.

69. Zhu, F., Kamiya, T., Fujiwara, T., et al. 2024. “A comparison of rice root microbial dynamics in organic and conventional paddy fields.” Microorganisms 13:41. 10.3390/microorganisms13010041.

