## Supporting Figures S1 to S8 and Supporting Tables S1-S2 for "Chemical warfare in the rice plant rhizosphere: Diverse *Pseudomonas* antagonize the green alga *Chlamydomonas*"

The following Supporting Information is available for this article:

Supporting Figure S1-S8: Pages 2 - 15

Supporting Table S1-S2: Pages 16 - 17

Supporting References: Page 18

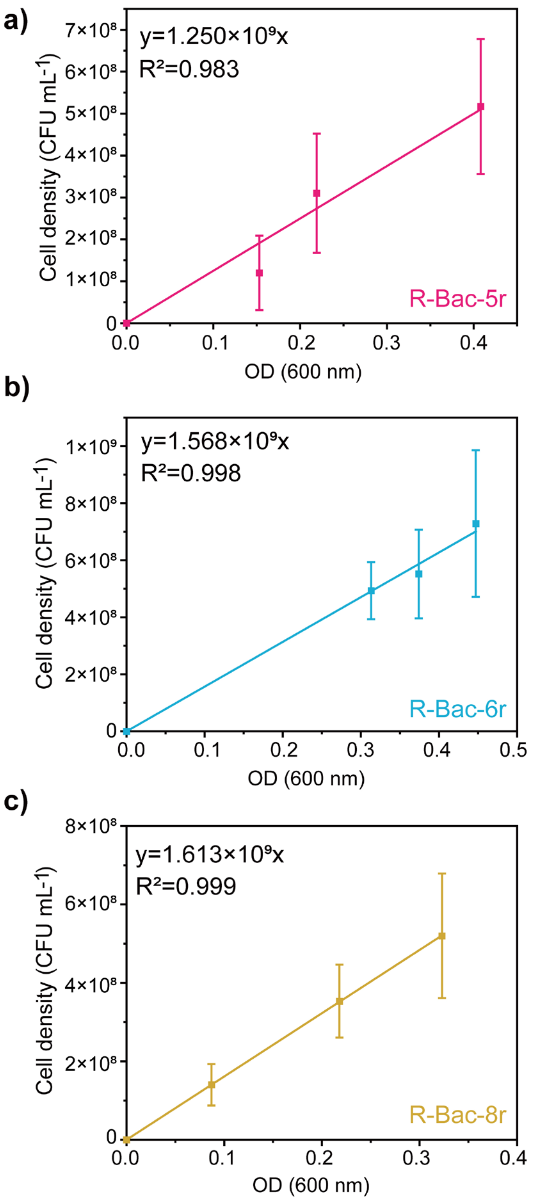


**FIGURE S1** | Calibration of optical density measurements for three antagonistic bacterial isolates. Correlation between $OD_{600}$ values measured in LB medium and bacterial cell density, determined as colony-forming units per milliliter (CFU $mL^{-1}$), for R-Bac-5r (a), R-Bac-6r (b), and R-Bac-8r (c), establishing the relationship between optical density and cell numbers.

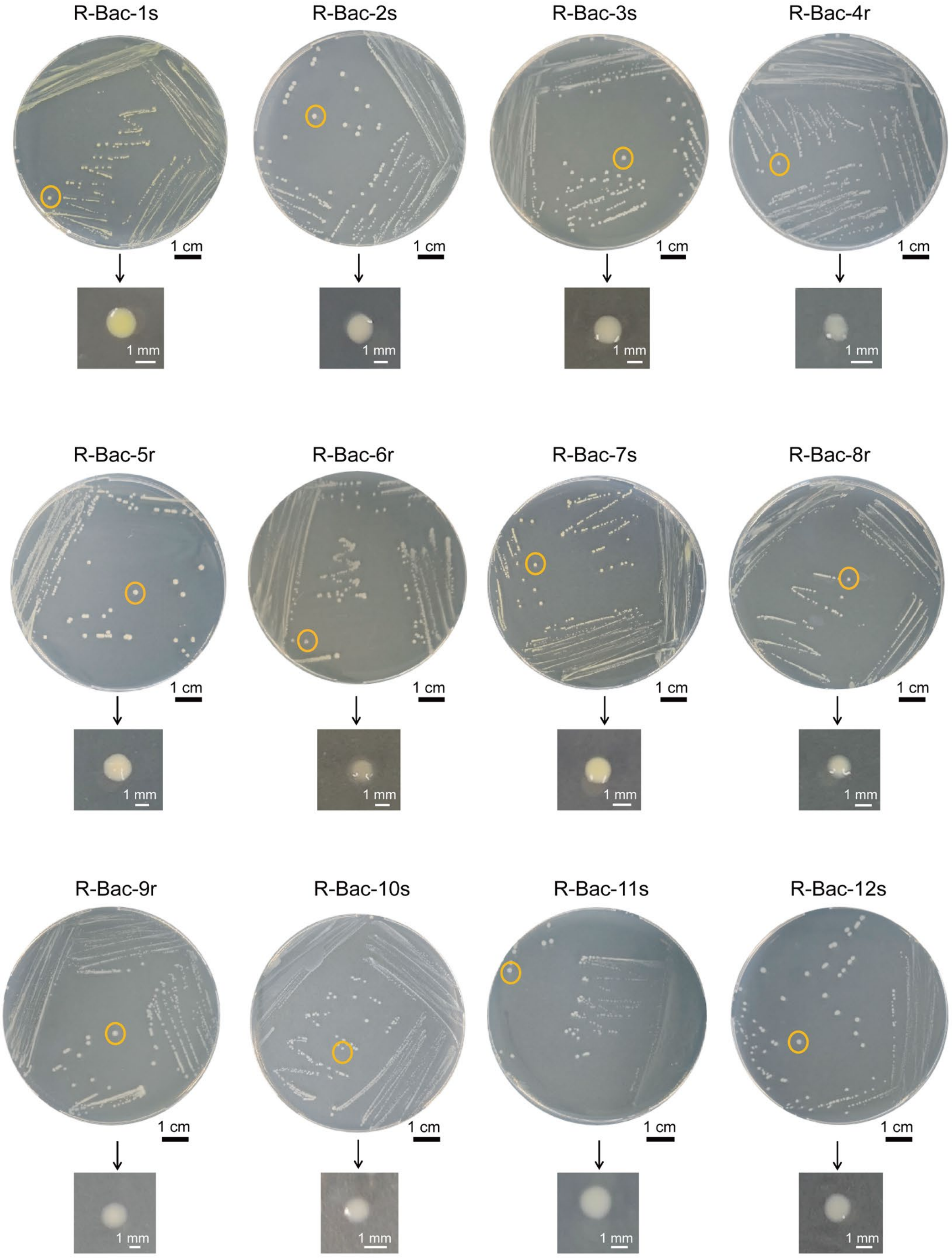


**FIGURE S2** | Colony morphology of 12 bacterial strains isolated from rice soil (s) or the rice rhizosphere (r). Strains were grown on TAP agar plates (Harris 1989). Representative colonies of each isolate are indicated by orange circles and shown individually beneath each plate.

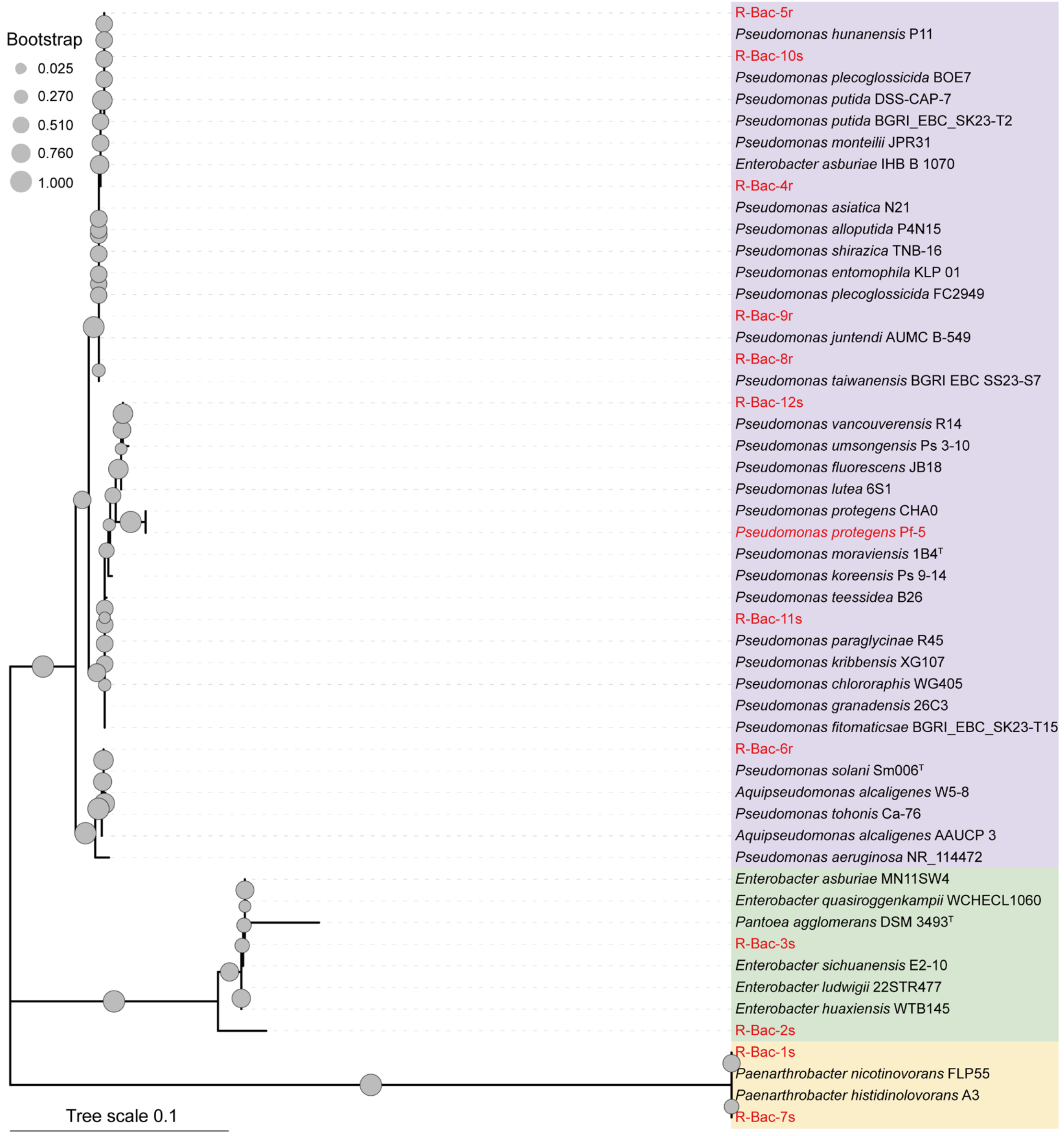


**FIGURE S3** | Phylogenetic relationships of bacterial isolates from the rice field based on *16S rRNA* gene sequences (Methods). Maximum likelihood phylogenetic analysis was performed to determine the evolutionary relationships among the 12 bacterial rice field strains (indicated in red color). The found genera *Pseudomonas*, *Enterobacter* along with the closely related *Pantoeae* as well as *Paenarthobacter* have been indicated in different background colours.

**R-Bac-1s**

AACGATGATCCCAGCTTGCTGGGGGATTAGTGGCGAACGGGTGAGTAACACGTGAGTAACCTGCCCTTGACT
CTGGGATAAGCCTGGGAAACTGGGTCTAATACCGGATATGACTCCTCATCGCATGGTGGGGGGTGGAAAGCT
TTTGTGGTTTTGGATGGACTCGCGGCCTATCAGCTTGTTGGTGGGGTAATGGCCTACCAAGGCGACGACGG
GTAGCCGGCCTGAGAGGGTGACCGGCCACACTGGGACTGAGACACGGCCCAGACTCCTACGGGAGGCAG
CAGTGGGGAATATTGCACAATGGGCGCAAGCCTGATGCAGCGACGCCGCGTGAGGGATGACGGCCTTCGG
GTTGTAAACCTCTTTCAGTAGGGAAGAAGCGTAAGTGACGGTACCTGCAGAAGAAGCGCCGGCTAACTACGT
GCCAGCAGCCGCGGTAATACGTAGGGCGCAAGCGTTATCCGGAATTATTGGGCGTAAAGAGCTCGTAGGCG
GTTTGTCGCGTCTGCTGTGAAAGACCGGGGCTCAACTCCGGTTCTGCAGTGGGTACGGGCAGACTAGAGTG
CAGTAGGGGAGACTGGAATTCCTGGTGTAGCGGTGAAATGCGCAGATATCAGGAGGAACACCGATGGCGAA
GGCAGGTCTCTGGGCTGTAACTGACGCTGAGGAGCGAAAGCATGGGGAGCGAACAGGATTAGATACCCTGG
TAGTCCATGCCGTAAACGTTGGGCACTAGGTGTGGGGGACATTCCACGTTTTCCGCGCCGTAGCTAACGCAT
TAAGTGCCCCGCCTGGGGAGTACGGCCGCAAGGCTAAAACTCAAAGGAATTGACGGGGGCCCGCACAAGC
GGCGGAGCATGCGGATTAATTCGATGCAACGCGAAGAACCTTACCAAGGCTTGACATGAACCGGAAAGACCT
GGAAACAGGTGCCCCGCTTGCGGTCGGTTTACAGGTGGTGCATGGTTGTCGTCAGCTCGTGTCGTGAGATG
TTGGGTTAAGTCCCGCAACGAGCGCAACCCTCGTTCTATGTTGCCAGCGGTTCGGCCGGGGACTCATAGGA
GACTGCCGGGGTCAACTCGGAGGAAGGTGGGGACGACGTCAAATCATCATGCCCCTTATGTCTTGGGCTTC
ACGCATGCTACAATGGCCGGTACAAAGGGTTGCGATACTGTGAGGTGGAGCTAATCCCAAAAAGCCGGTCTC
AGTTCGGATTGGGGTCTGCAACTCGACCCCATGAAGTCGGAGTCGCTAGTAATCGCAGATCAGCAACGCTGC
GGTGAATACGTTCCCGGGCCTTGTACACACCGCCCGTCAAGTCACGAAAGTTGGTAACACCCGAAGCCGGT
GGCCTAACC

**R-Bac-2s**

TGGCTCAGAGTTTGATCCTGGCTCAGAGTTTGATCCTGGCTCAGAGTTTGATCCTGGCTCAGAGTTTGATCCT
GGCTCACAGTTTGATCCTGGCTCATACTTTGATCCTGGCTCAAACATTGAGGGGGACCTTCAGGCCTCTTGC
CATCAGAAGTGCCCTCATGGGATTAACTANTAGGTGGGGTAACGGCTCACCTAGGCGACGATCCCTAGCTGG
TCTGAGAGGATGACCAGCCACACTGGAACTGAGACACGGTCCAGACTCCTACGGGAGGCAGCAGTGGGGA
ATATTGCACAATGGGCGCAAGCCTGATGCAGCCATGCCGCGTGTATGAAGAAGGCCTTCGGGTTGTAAAGTA
CTTTCAGCGGGGAGGAAGGTGTTGTGGTTAATAACCGCAGCAATTGACGTTACCCGCAGAAGAAGCACCGG
CTAACTCCGTGCCAGCAGCCGCGGTAATACGGAGGGTGCAAGCGTTAATCGGAATTACTGGGCGTAAAGCG
CACGCAGGCGGTCTGTCAAGTCGGATGTGAAATCCCCGGGCTCAACCTGGGAACTGCATTCGAAACTGGCA
GGCTAGAGTCTTGTAGAGGGGGGTAGAATTCCAGGTGTAGCGGTGAAATGCGTAGAGATCTGGAGGAATACC
GGTGGCGAAGGCGGCCCCCTGGACAAAGACTGACGCTCAGGTGCGAAAGCGTGGGGAGCAAACAGGATTA
GATACCCTGGTAGTCCACGCCGTAAACGATGTCGACTTGGAGGTTGTGCCCTTGAGGCGTGGCTTCCGGAG
CTAACGCGTTAAGTCGACCGCCTGGGGAGTACGGCCGCAAGGTTAAAACTCAAATGAATTGACGGGGGCCC
GCACAAGCGGTGGAGCATGTGGTTTAATTCGATGCAACGCGAAGAACCTTACCTACTCTTGACATCCAGAGA
ACTTTCCAGAGATGGATTGGTGCCTTCGGGAACTCTGAGACAGGTGCTGCATGGCTGTCGTCAGCTCGTGTT
GTGAAATGTTGGGTTAAGTCCCGCAACGAGCGCAACCCTTATCCTTTGTTGCCAGCGGTCCGGCCGGGAAC
TCAAAGGAGACTGCCAGTGATAAACTGGAGGAAGGTGGGGATGACGTCAAGTCATCATGGCCCTTACGAGTA
GGGCTACACACGTGCTACAATGGCGCATACAAAGAGAAGCGA

**R-Bac-3s**

CGGCTCACCTAGGCGACGATCCCTAGCTGGTCTGAGAGGATGACCAGCCACACTGGAACTGAGACACGGTC
CAGACTCCTACGGGAGGCAGCAGTGGGGAATATTGCACAATGGGCGCAAGCCTGATGCAGCCATGCCGCGT

GTATGAAGAAGGCCTTCGGAGTTGTAATAGTACTTTCAGCGGGTGAGGAAGGTGTTGAGGTTAATAACCTCAT
CAATTGACGTTACCCGCANAAGAAGCACCGGCTAACTCCGTGCCANCAGCCGCGGTAATACGGAGGGTGCA
AGCGTTAATCGNAATTACTGGGCGTAAANCNCACGCAGGCGGTCTGTCAAGTCNGATGTGAAATCCCCGGGC
TCAACCTGGGAACTGCATTCGAAACTGGCAGGCTAGAGTCTTGTAGAGGGGGGTAGAATTCCAGGTGTAGC
GGTGAAATGCGTANAGATCTGGAGGAATACCGGTGGCGAAGGCGGCCCCCTGGACAAAGACTGACNCTCAG
GTGCGAAAGCGTGGGGAGCAAACAGGATTAGATACCCTGGTAGTCCACGCCGTAAACGATGTCNACTTGNA
GGTTGTGCCCTTGAGGCGTGGCTTCCGGAGCTAACGCGTTAAGTCGACCGCCTGGGGAGTACGGCCGCAA
GGTTAAAACTCAAATGAATTGACGGGGGCCCGCACAAGCGGTGGAGCATGTGGTTTAATTCGATGCAACGCG
AAGAACCTTACCTACTCTTGACATCCAGAGAACTTTCCAGAGATGGATTGGTGCCTTCGGGAACTCTGAGACA
GGTGCTGCATGGCTGTCGTCAGCTCGTGTTGTGAAATGTTGGGTTAAGTCCCGCAACGAGCGCAACCCTTAT
CCTTTGTTGCCAGCGGTTCGGCCGGGAACTCAAAGGAGACTGCCAGTGATAAACTGGAGGAAGGTGGGGAT
GACGTCAAGTCATCATGGCCCTTACGAGTAGGGCTACACACGTGCTACAATGGCGCATACAAAGAGAAGCGA
CCTCGCGAGAGCAAGCGGACCTCATAAAGTGCGT

**R-Bac-4r**

GACAACGTTTCGAAAGGAACGCTAATACCGCATACGTCCTACGGGAGAAAGCAGGGGACCTTCGGGCCTTG
CGCTATCAGATGAGCCTAGGTCGGATTAGCTAGTTGGTGGGGTAATGGCTCACCAAGGCGACGATCCGTAAC
TGGTCTGAGAGGATGATCAGTCACACTGGAACTGAGACACGGTCCAGACTCCTACGGGAGGCAGCAGTGGG
GAATATTGGACAATGGGCGAAAGCCTGATCCAGCCATGCCGCGTGTGTGAAGAAGGTCTTCGGATTGTAAAG
CACTTTAAGTTGGGAGGAAGGGCAGTAAGNTAATACCTTGCTGTTTTGACGTTACCGACAGAATAAGCACCGG
CTAACTCTGTGCCAGCAGCCGCGGTAATACAGAGGGTGCAAGCGTTAATCGGAATTACTGGGCGTAAAGCGC
GCGTAGGTGGTTCGTTAAGTTGGATGTGAAAGCCCCGGGCTCAACCTGGGAACTGCATCCAAAACTGGCGA
GCTAGAGTACGGTAGAGGGTGGTGGAATTTCCTGTGTAGCGGTGAAATGCGTAGATATAGGAAGGAACACCA
GTGGCGAAGGCGACCACCTGGACTGATACTGACACTGAGGTGCGAAAGCGTGGGGAGCAAACAGGATTAG
ATACCCTGGTAGTCCACGCCGTAAACGATGTCAACTAGCCGTTGGAATCCTTGAGATTTTAGTGGCGCAGCTA
ACGCATTAAGTTGACCGCCTGGGGAGTACGGCCGCAAGGTTAAAACTCAAATGAATTGACGGGGGCCCGCA
CAAGCGGTGGAGCATGTGGTTTAATTCGAAGCAACGCGAAGAACCTTACCAGGCCTTGACATGCAGAGAACT
TTCCAGAGATGGATTGGTGCCTTCGGGAACTCTGACACAGGTGCTGCATGGCTGTCGTCAGCTCGTGTCGT
GAGATGTTGGGTTAAGTCCCGTAACGAGCGCAACCCTTGTCCTTAGTTACCAGCACGTTATGGTGGGCACTC
TAAGGAGACTGCCGGTGACAAACCGGAGGAAGGTGGGGATGACGTCAAGTCATCATGGCCCTTACGGCCTG
GGCTACACACGTGCTACAATGGTCGGTACAGAGGGTTGCCAAGCCGCGAGGTGGAGCTAATCTCACAAAAC
CGATCGTAGTCCGGATCGCAGTCTGCAACTCGACTGCGTGAAGTCGGAATCGCTAGTAATCGCGAATCAGAA
TGTCGCGGTGAATACGTTCCC

**R-Bac-5r**

TAATGCCTAGGAATCTGCCTGGTAGTGGGGGACAACGTTTCGAAAGGAACGCTAATACCGCATACGTCCTACG
GGAGAAAGCAGGGGACCTTCGGGCCTTGCGCTATCAGATGAGCCTAGGTCGGATTAGCTAGTTGGTGGGGT
AATGGCTCACCAAGGCGACGATCCGTAACTGGTCTGAGAGGATGATCAGTCACACTGGAACTGAGACACGG
TCCAGACTCCTACGGGAGGCAGCAGTGGGGAATATTGGACAATGGGCGAAAGCCTGATCCAGCCATGCCGC
GTGTGTGAAGAAGGTCTTCGGATTGTAAAGCACTTTAAGTTGGGAGGAAGGGCAGTAAGCTAATACCTTGCT
GTTTTGACGTTACCGACAGAATAAGCACCGGCTAACTCTGTGCCAGCAGCCGCGGTAATACAGAGGGTGCAA
GCGTTAATCGGAATTACTGGGCGTAAAGCGCGCGTAGGTGGTTTGTTAAGTTGGATGTGAAAGCCCCGGGCT
CAACCTGGGAACTGCATCCAAAACTGGCAAGCTAGAGTACGGTAGAGGGTGGTGGAATTTCCTGTGTAGCG
GTGAAATGCGTAGATATAGGAAGGAACACCAGTGGCGAAGGCGACCACCTGGACTGATACTGACACTGAGGT

GCGAAAGCGTGGGGAGCAAACAGGATTAGATACCCTGGTAGTCCACGCCGTAAACGATGTCAACTAGCCGTT
GGAATCCTTGAGATTTTAGTGGCGCAGCTAACGCATTAAGTTGACCGCCTGGGGAGTACGGCCGCAAGGTTA
AAACTCAAATGAATTGACGGGGGCCCGCACAAGCGGTGGAGCATGTGGTTTAATTCGAAGCAACGCGAAGA
ACCTTACCAGGCCTTGACATGCAGAGAACTTTCCAGAGATGGATTGGTGCCTTCGGGAACTCTGACACAGGT
GCTGCATGGCTGTCGTCAGCTCGTGTCGTGAGATGTTGGGTTAAGTCCCGTAACGAGCGCAACCCTTGTCCT
TAGTTACCAGCACGTTATGGTGGGCACTCTAAGGAGACTGCCGGTGACAAACCGGAGGAAGGTGGGGATGA
CGTCAAGTCATCATGGCCCTTACGGCCTGGGCTACACACGTGCTACAATGGTCGGTACAGAGGGTTGCCAAG
CCGCGAGGTGGAGCTAATCTCACAAAACCGATCGTAGTCCGGATCGCAGTCTGCAACTCGACTGCGTGAAG
TCGGAATCGCTAGTAATCGCGAATCAGAATGTCGCGGTGAATACGTTCCCGGGCCTTGTACACACCGCCCGT
CACACCATGGGAGTGGGTGCACCAGAAGTAGCTAGTCTAACCTTCGG

**R-Bac-6r**

TGGAGCTTGCTCGATGATTCAGCGGCAGGACGGGTGAGTAATGCCTAGGAATCTGCCTGGTAGTGGGGGAC
AACGTTTCGAAAGGAACGCTAATACCGCATACGTCCTACGGGAGAAAGCAGGGGACCTTCGGGCCTTGCGC
TATCAGATGAGCCTAGGTCGGATTAGCTAGTTGGTGAGGTAAAGGCTCACCAAGGCGACGATCCGTAACTGG
TCTGAGAGGATGATCAGTCACACTGGAACTGAGACACGGTCCAGACTCCTACGGGAGGCAGCAGTGGGGAA
TATTGGACAATGGGCGAAAGCCTGATCCAGCCATGCCGCGTGTGTGAAGAAGGTCTTCGGATTGTAAAGCAC
TTTAAGTTGGGAGGAAGGGCAGTAAGTTAATACCTTGCTGTTTTGACGTTACCGACAGAATAAGCACCGGCTA
ACTTCGTGCCAGCAGCCGCGGTAATACGAAGGGTGCAAGCGTTAATCGGAATTACTGGGCGTAAAGCGCGC
GTAGGTGGTTCAGCAAGTTGGATGTGAAAGCCCCGGGCTCAACCTGGGAACTGCATCCAAAACTACTGAGC
TAGAGTACGGTAGAGGGTGGTGGAATTTCCTGTGTAGCGGTGAAATGCGTAGATATAGGAAGGAACACCAGT
GGCGAAGGCGACCACCTGGACTGATACTGACACTGAGGTGCGAAAGCGTGGGGAGCAAACAGGATTAGATA
CCCTGGTAGTCCACGCCGTAAACGATGTCGACTAGCCGTTGGGATCCTTGAGATCTTAGTGGCGAAGCTAAC
GCGATAAGTCGACCGCCTGGGGAGTACGGCCGCAAGGTTAAAACTCAAATGAATTGACGGGGGCCCGCACA
AGCGGTGGAGCATGTGGTTTAATTCGAAGCAACGCGAAGAACCTTACCTGGCCTTGACATGCTGAGAACTTT
CCAGAGATGGATTGGTGCCTTCGGGAACTCAGACACAGGTGCTGCATGGCTGTCGTCAGCTCGTGTCGTGA
GATGTTGGGTTAAGTCCCGTAACGAGCGCAACCCTTGTCCTTAGTTACCAGCACGTTATGGTGGGCACTCTAA
GGAGACTGCCGGTGACAAACCGGAGGAAGGTGGGGATGACGTCAAGTCATCATGGCCCTTACGGCCAGGG
CTACACACGTGCTACAATGGTCGGTACAAAGGGTTGCCAAGCCGCGAGGTGGAGCTAATCCCATAAAACCGA
TCGTAGTCCGGATCGCAGTCTGCAACTCGACTGCGTGAAGTCGGAATCGCTAGTAATCGTGAATCAGAATGT
CACGGTGAATACGTTCCCGGGCCTGTACACACCGCCCGTCACACCATGGGAGTGGGCTGCTCCAGAAGTAG
C

**R-Bac-7s**

CGAACGATGATCCCAGCTTGCTGGGGGATAGTGGCGAACGGGTGAGTAACACGTGAGTAACCTGCCCTTGA
CTCTGGGATAAGCCTGGGAAACTGGGTCTAATACCGGATATGACTCCTCATCGCATGGTGGGGGGTGGAAAG
CTTTTGTGGTTTTGGATGGACTCGCGGCCTATCAGCTTGTTGGTGGGGTAATGGCCTACCAAGGCGACGACG
GGTAGCCGGCCTGAGAGGGTGACCGGCCACACTGGGACTGAGACACGGCCCAGACTCCTACGGGAGGCA
GCAGTGGGGAATATTGCACAATGGGCGCAAGCCTGATGCAGCGACGCCGCGTGAGGGATGACGGCCTTCG
GGTTGTAAACCTCTTTCAGTAGGGAAGAAGCGTAAGTGACGGTACCTGCAGAAGAAGCGCCGGCTAACTACG
TGCCAGCAGCCGCGGTAATACGTAGGGCGCAAGCGTTATCCGGAATTATTGGGCGTAAAGAGCTCGTAGGCG
GTTTGTCGCGTCTGCTGTGAAAGACCGGGGCTCAACTCCGGTTCTGCAGTGGGTACGGGCAGACTAGAGTG
CAGTAGGGGAGACTGGAATTCCTGGTGTAGCGGTGAAATGCGCAGATATCAGGAGGAACACCGATGGCGAA
GGCAGGTCTCTGGGCTGTAACTGACGCTGAGGAGCGAAAGCATGGGGAGCGAACAGGATTAGATACCCTGG

TAGTCCATGCCGTAAACGTTGGGCACTAGGTGTGGGGGACATTCCACGTTTTCCGCGCCGTAGCTAACGCAT
TAAGTGCCCCGCCTGGGGAGTACGGCCGCAAGGCTAAAACTCAAAGGAATTGACGGGGGCCCGCACAAGC
GGCGGAGCATGCGGATTAATTCGATGCAACGCGAAGAACCTTACCAAGGCTTGACATGAACCGGAAAGACCT
GGAAACAGGTGCCCCGCTTGCGGTCGGTTTACAGGTGGTGCATGGTTGTCGTCAGCTCGTGTCGTGAGATG
TTGGGTTAAGTCCCGCAACGAGCGCAACCCTCGTTCTATGTTGCCAGCGGTTCGGCCGGGGACTCATAGGA
GACTGCCGGGGTCAACTCGGAGGAAGGTGGGGACGACGTCAAATCATCATGCCCCTTATGTCTTGGGCTTC
ACGCATGCTACAATGGCCGGTACAAAGGGTTGCGATACTGTGAGGTGGAGCTAATCCCAAAAAGCCGGTCTC
AGTTCGGATTGGGGTCTGCAACTCGACCCCATGAAGTCGGAGTCGCTAGTAATCGCAGATCAGCAACGCTGC
GGTGAATACGTTCCCGGGCCTTGTACACACCGCCCGTCAAGTCACGAAAGTTGGTAACACCCGAAGCCGGT
GGCCTAACCCTTGTGGGG

**R-Bac-8r**

CGAGCGGATGACGGGAGCTTGCTCCTTGATTCAGCGGCGGACGGGTGAGTAATGCCTAGGAATCTGCCTGG
TAGTGGGGGACAACGTTTCGAAAGGAACGCTAATACCGCATACGTCCTACGGGAGAAAGCAGGGGACCTTC
GGGCCTTGCGCTATCAGATGAGCCTAGGTCGGATTAGCTAGTTGGTGAGGTAATGGCTCACCAAGGCGACGA
TCCGTAACTGGTCTGAGAGGATGATCAGTCACACTGGAACTGAGACACGGTCCAGACTCCTACGGGAGGCA
GCAGTGGGGAATATTGGACAATGGGCGAAAGCCTGATCCAGCCATGCCGCGTGTGTGAAGAAGGTCTTCGG
ATTGTAAAGCACTTTAAGTTGGGAGGAAGGGCAGTAAGTTAATACCTTGCTGTTTTGACGTTACCGACAGAATA
AGCACCGGCTAACTCTGTGCCAGCAGCCGCGGTAATACAGAGGGTGCAAGCGTTAATCGGAATTACTGGGC
GTAAAGCGCGCGTAGGTGGTTCGTTAAGTTGGATGTGAAAGCCCCGGGCTCAACCTGGGAACTGCATCCAA
AACTGGCGAGCTAGAGTACGGTAGAGGGTGGTGGAATTTCCTGTGTAGCGGTGAAATGCGTAGATATAGGAA
GGAACACCAGTGGCGAAGGCGACCACCTGGACTGATACTGACACTGAGGTGCGAAAGCGTGGGGAGCAAA
CAGGATTAGATACCCTGGTAGTCCACGCCGTAAACGATGTCAACTAGCCGTTGGAATCCTTGAGATTTTAGTG
GCGCAGCTAACGCATTAAGTTGACCGCCTGGGGAGTACGGCCGCAAGGTTAAAACTCAAATGAATTGACGGG
GGCCCGCACAAGCGGTGGAGCATGTGGTTTAATTCGAAGCAACGCGAAGAACCTTACCAGGCCTTGACATG
CAGAGAACTTTCCAGAGATGGATTGGTGCCTTCGGGAACTCTGACACAGGTGCTGCATGGCTGTCGTCAGC
TCGTGTCGTGAGATGTTGGGTTAAGTCCCGTAACGAGCGCAACCCTTGTCCTTAGTTACCAGCACGTTATGGT
GGGCACTCTAAGGAGACTGCCGGTGACAAACCGGAGGAAGGTGGGGATGACGTCAAGTCATCATGGCCCTT
ACGGCCTGGGCTACACACGTGCTACAATGGTCGGTACAGAGGGTTGCCAAGCCGCGAGGTGGAGCTAATCT
CACAAAACCGATCGTAGTCCGGATCGCAGTCTGCAACTCGACTGCGTGAAGTCGGAATCGCTAGTAATCGCG
AATCAGAATGTCGCGGTGAATACGTTCCCGGGCCTTGTACACACCGCCCGTCACACCATGGGAGTGGGTGC
ACCAGAAGTAGCTAGTCTAACCTTCGGGAGGA

**R-Bac-9r**

ATGCCTAGGAATCTGCCTGGTAGTGGGGGACAACGTTTCGAAAGGAACGCTAATACCGCATACGTCCTACGG
GAGAAAGCAGGGGACCTTCGGGCCTTGCGCTATCAGATGAGCCTAGGTCGGATTAGCTAGTTGGTGAGGTAA
TGGCTCACCAAGGCGACGATCCGTAACTGGTCTGAGAGGATGATCAGTCACACTGGAACTGAGACACGGTC
CAGACTCCTACGGGAGGCAGCAGTGGGGAATATTGGACAATGGGCGAAAGCCTGATCCAGCCATGCCGCGT
GTGTGAAGAAGGTCTTCGGATTGTAAAGCACTTTAAGTTGGGAGGAAGGGCAGTAAGTTAATACCTTGCTGTT
TTGACGTTACCGACAGAATAAGCACCGGCTAACTCTGTGCCAGCAGCCGCGGTAATACAGAGGGTGCAAGC
GTTAATCGGAATTACTGGGCGTAAAGCGCGCGTAGGTGGTTCGTTAAGTTGGATGTGAAAGCCCCGGGCTCA
ACCTGGGAACTGCATCCAAAACTGGCGAGCTAGAGTACGGTAGAGGGTGGGTGGAATTTCCTGTGTAGCGG
TGAAATGCGTAGATATAGGAAGGAACACCAGTGGCGAAGGCGACCACCTGGACTGATACTGACACTGAGGTG
CGAAAGCGTGGGGAGCAAACAGGATTAGATACCCTGGTAGTCCACGCCGTAAACGATGTCAACTAGCCGTTG

GAATCCTTGAGATTTTAGTGGCGCAGCTAACGCATTAAGTTGACCGCCTGGGGAGTACGGCCGCAAGGTTAA
AACTCAAATGAATTGACGGGGGCCCGCACAAGCGGTGGAGCATGTGGTTTAATTCGAAGCAACGCGAAGAA
CCTTACCAGGCCTTGACATGCAGAGAACTTTCCAGAGATGGATTGGTGCCTTCGGGAACTCTGACACAGGTG
CTGCATGGCTGTCGTCAGCTCGTGTCGTGAGATGTTGGGTTAAGTCCCGTAACGAGCGCAACCCTTGTCCTT
AGTTACCAGCACGTTATGGTGGGCACTCTAAGGAGACTGCCGGTGACAAACCGGAGGAAGGTGGGGATGAC
GTCAAGTCATCATGGCCCTTACGGCCTGGGCTACACACGTGCTACAATGGTCGGTACAGAGGGTTGCCAAGC
CGCGAGGTGGAGCTAATCTCACAAAACCGATCGTAGTCCGGATCGCAGTCTGCAACTCGACTGCGTGAAGT
CGGAATCGCTAGTAATCGCGAATCAGAATGTCGCGGTGAATACGTTCCCGGGCCTTGTACACACCGCCCGTC
ACACCATGGGAGTGGGTGCACCAGAAGTAGCTAGTCTAACCTTCG

**R-Bac-10s**

TAGTGGGGGACAACGTTTCGAAAGGAACGCTAATACCGCATACGTCCTACGGGAGAAAGCAGGGGACCTTC
GGGCCTTGCGCTATCAGATGAGCCTAGGTCGGATTAGCTAGTTGGTGGGGTAATGGCTCACCAAGGCGACG
ATCCGTAACTGGTCTGAGAGGATGATCAGTCACACTGGAACTGAGACACGGTCCAGACTCCTACGGGAGGC
AGCAGTGGGGAATATTGGACAATGGGCGAAAGCCTGATCCAGCCATGCCGCGTGTGTGAAGAAGGTCTTCG
GATTGTAAAGCACTTTAAGTTGGGAGGAAGGGCAGTAAGCTAATACCTTGCTGTTTTGACGTTACCGACAGAA
TAAGCACCGGCTAACTCTGTGCCAGCAGCCGCGGTAATACAGAGGGTGCAAGCGTTAATCGGAATTACTGGG
CGTAAAGCGCGCGTAGGTGGTTTGTTAAGTTGGATGTGAAAGCCCCGGGCTCAACCTGGGAACTGCATCCA
AAACTGGCAAGCTAGAGTACGGTAGAGGGTGGTGGAATTTCCTGTGTAGCGGTGAAATGCGTAGATATAGGA
AGGAACACCAGTGGCGAAGGCGACCACCTGGACTGATACTGACACTGAGGTGCGAAAGCGTGGGGAGCAA
ACAGGATTAGATACCCTGGTAGTCCACGCCGTAAACGATGTCAACTAGCCGTTGGAATCCTTGAGATTTTAGT
GGCGCAGCTAACGCATTAAGTTGACCGCCTGGGGAGTACGGCCGCAAGGTTAAAACTCAAATGAATTGACGG
GGGCCCGCACAAGCGGTGGAGCATGTGGTTTAATTCGAAGCAACGCGAAGAACCTTACCAGGCCTTGACAT
GCAGAGAACTTTCCAGAGATGGATTGGTGCCTTCGGGAACTCTGACACAGGTGCTGCATGGCTGTCGTCAG
CTCGTGTCGTGAGATGTTGGGTTAAGTCCCGTAACGAGCGCAACCCTTGTCCTTAGTTACCAGCACGTTATG
GTGGGCACTCTAAGGAGACTGCCGGTGACAAACCGGAGGAAGGTGGGGATGACGTCAAGTCATCATGGCC
CTTACGGCCTGGGCTACACACGTGCTACAATGGTCGGTACAGAGGGTTGCCAAGCCGCGAGGTGGAGCTAA
TCTCACAAAACCGATCGTAGTCCGGATCGCAGTCTGCAACTCGACTGCGTGAAGTCGGAATCGCTAGTAATC
GCGAATCAGAATGTCGCGGTGAATACGTTCCC

**R-Bac-11s**

AGCGGATGAAGGGAGCTTGCTCCTGGATTCAGCGGCGGACGGGTGAGTAATGCCTAGGAATCTGCCTGGTA
GTGGGGGACAACGTTTCGAAAGGAACGCTAATACCGCATACGTCCTACGGGAGAAAGCAGGGGACCTTCGG
GCCTTGCGCTATCAGATGAGCCTAGGTCGGATTAGCTAGTTGGTGAGGTAATGGCTCACCAAGGCGACGATC
CGTAACTGGTCTGAGAGGATGATCAGTCACACTGGAACTGAGACACGGTCCAGACTCCTACGGGAGGCAGC
AGTGGGGAATATTGGACAATGGGCGAAAGCCTGATCCAGCCATGCCGCGTGTGTGAAGAAGGTCTTCGGATT
GTAAAGCACTTTAAGTTGGGAGGAAGGGTTGTAGATTAATACTCTGCAATTTTGACGTTACCGACAGAATAAGC
ACCGGCTAACTCTGTGCCAGCAGCCGCGGTAATACAGAGGGTGCAAGCGTTAATCGGAATTACTGGGCGTAA
AGCGCGCGTAGGTGGTTCGTTAAGTTGGATGTGAAATCCCCGGGCTCAACCTGGGAACTGCATCCAAAACTG
GCGAGCTAGAGTATGGTAGAGGGTGGTGGAATTTCCTGTGTAGCGGTGAAATGCGTAGATATAGGAAGGAAC
ACCAGTGGCGAAGGCGACCACCTGGACTGATACTGACACTGAGGTGCGAAAGCGTGGGGAGCAAACAGGA
TTAGATACCCTGGTAGTCCACGCCGTAAACGATGTCAACTAGCCGTTGGGAGCCTTGAGCTCTTAGTGGCGC
AGCTAACGCATTAAGTTGACCGCCTGGGGAGTACGGCCGCAAGGTTAAAACTCAAATGAATTGACGGGGGCC
CGCACAAGCGGTGGAGCATGTGGTTTAATTCGAAGCAACGCGAAGAACCTTACCAGGCCTTGACATCCAATG

AACTTTCCAGAGATGGATTGGTGCCTTCGGGAACATTGAGACAGGTGCTGCATGGCTGTCGTCAGCTCGTGT
CGTGAGATGTTGGGTTAAGTCCCGTAACGAGCGCAACCCTTGTCCTTAGTTACCAGCACGTAATGGTGGGCA
CTCTAAGGAGACTGCCGGTGACAAACCGGAGGAAGGTGGGGATGACGTCAAGTCATCATGGCCCTTACGGC
CTGGGCTACACACGTGCTACAATGGTCGGTACAAAGGGTTGCCAAGCCGCGAGGTGGAGCTAATCCCATAAA
ACCGATCGTAGTCCGGATCGCAGTCTGCAACTCGACTGCGTGAAGTCGGAATCGCTAGTAATCGCGAATCAG
AATGTCGCGGTGAATACGTTCCCGGGCCTTGTACACACCGCCCGTCACACCATGGGAGTGGGTGCACCAGA
AGTAGCTAGTCTAACCTTCGGGAG

**R-Bac-12s**
AGCGGATGAGAGAGCTTGCTCTCTGATTCAGCGGCGGACGGGTGAGTAATGCCTAGGAATCTGCCTGGTAG
TGGGGGACAACGTCTCGAAAGGGACGCTAATACCGCATACGTCCTACGGGAGAAAGCAGGGGACCTTCGGG
CCTTGCGCTATCAGATGAGCCTAGGTCGGATTAGCTAGTTGGTGAGGTAATGGCTCACCAAGGCGACGATCC
GTAACTGGTCTGAGAGGATGATCAGTCACACTGGAACTGAGACACGGTCCAGACTCCTACGGGAGGCAGCA
GTGGGGAATATTGGACAATGGGCGAAAGCCTGATCCAGCCATGCCGCGTGTGTGAAGAAGGTCTTCGGATT
GTAAAGCACTTTAAGTTGGGAGGAAGGGTTGTAGATTAATACTCTGCAATTTTGACGTTACCGACAGAATAAGC
ACCGGCTAACTCTGTGCCAGCAGCCGCGGTAATACAGAGGGTGCAAGCGTTAATCGGAATTACTGGGCGTAA
AGCGCGCGTAGGTGGTTTGTTAAGTTGGATGTGAAAGCCCCGGGCTCAACCTGGGAACTGCATTCAAAACT
GACAAGCTAGAGTATGGTAGAGGGTGGTGGAATTTCCTGTGTAGCGGTGAAATGCGTAGATATAGGAAGGAA
CACCAGTGGCGAAGGCGACCACCTGGACTGATACTGACACTGAGGTGCGAAAGCGTGGGGAGCAAACAGG
ATTAGATACCCTGGTAGTCCACGCCGTAAACGATGTCAACTAGCCGTTGGGAGCCTTGAGCTCTTAGTGGCG
CAGCTAACGCATTAAGTTGACCGCCTGGGGAGTACGGCCGCAAGGTTAAAACTCAAATGAATTGACGGGGGC
CCGCACAAGCGGTGGAGCATGTGGTTTAATTCGAAGCAACGCGAAGAACCTTACCAGGCCTTGACATCCAAT
GAACTTTCCAGAGATGGATTGGTGCCTTCGGGAACATTGAGACAGGTGCTGCATGGCTGTCGTCAGCTCGT
GTCGTGAGATGTTGGGTTAAGTCCCGTAACGAGCGCAACCCTTGTCCTTAGTTACCAGCACGTAATGGTGGG
CACTCTAAGGAGACTGCCGGTGACAAACCGGAGGAAGGTGGGGATGACGTCAAGTCATCATGGCCCTTACG
GCCTGGGCTACACACGTGCTACAATGGTCGGTACAGAGGGTTGCCAAGCCGCGAGGTGGAGCTAATCCCAT
AAAACCGATCGTAGTCCGGATCGCAGTCTGCAACTCGACTGCGTGAAGTCGGAATCGCTAGTAATCGCGAAT
CAGAATGTCGCGGTGAATACGTTCCCGGGCCTTGTACACACCGCCCGTCACACCATGGGAGTGGGTGCACC
AGAAGTAGCTAGTCTAACCTTCGGGAGGA

**FIGURE S4** | Amplified *16S rRNA* gene sequences (Methods) of 12 bacterial isolates from the rice field soil (s) or the rice rhizosphere (r), presented in the 5′–3′ direction.

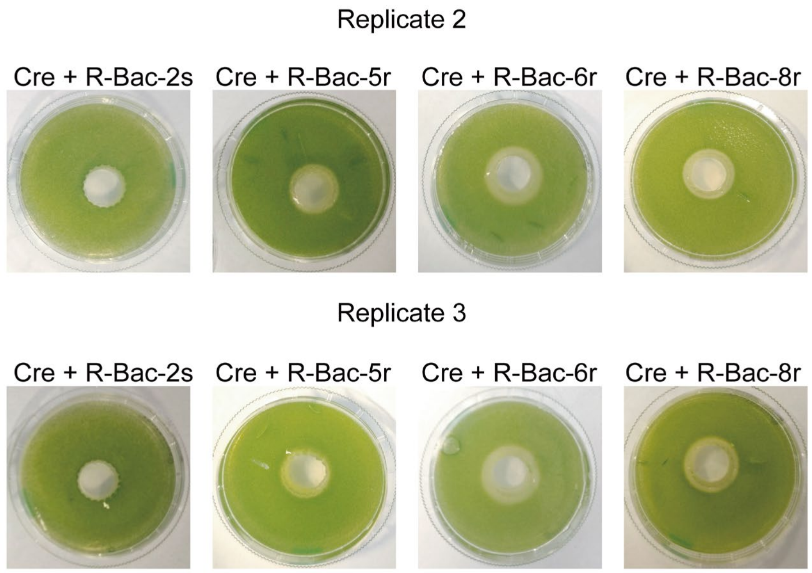


**FIGURE S5** | Antagonistic rice rhizosphere isolates R-Bac-5r, R-Bac-6r, and R-Bac-8r compared to the commensal R-Bac-2s. Growth inhibition agar plate assays of *Chlamydomonas reinhardtii* (Cre) with bacterial isolates from the rice field. Bacteria were inoculated on a circular area in the middle of the agar plate that was covered fully by an algal top layer (Methods). Images were taken after five days of growth. Isolate R-Bac-2s showed no inhibition zone (commensal bacterium). Bacteria R-Bac-5r, R-Bac-6r, and R-Bac-8r exhibited inhibitory effects on Cre growth. Replicates 2 and 3 are presented, replicate 1 is shown in Figure 1d.

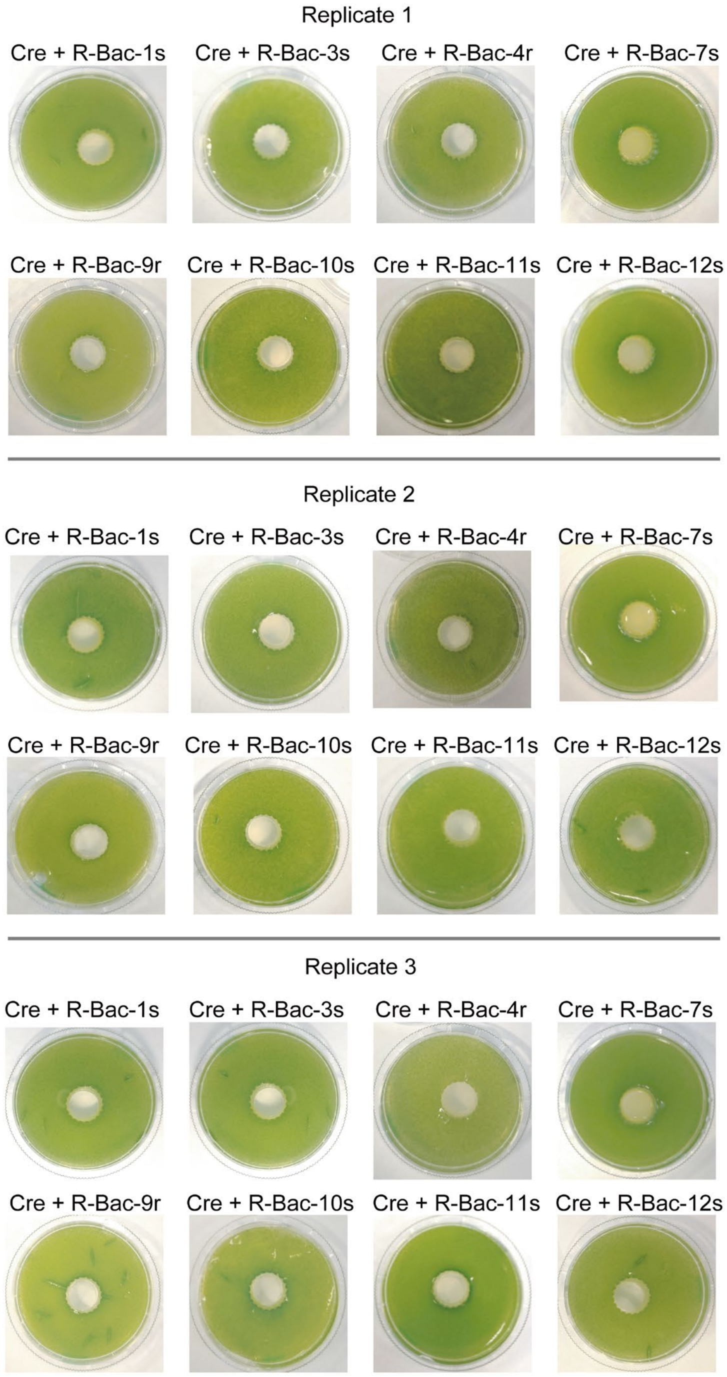


**FIGURE S6** | Screening of rice field bacterial isolates against *C. reinhardtii* (Cre). Commensal bacteria having no effect on the growth of Cre are shown, including R-Bac 1s, 3s, 4r, 7s, 9r, 10s, 11s and 12s. Assays were conducted as described in Figure S5.

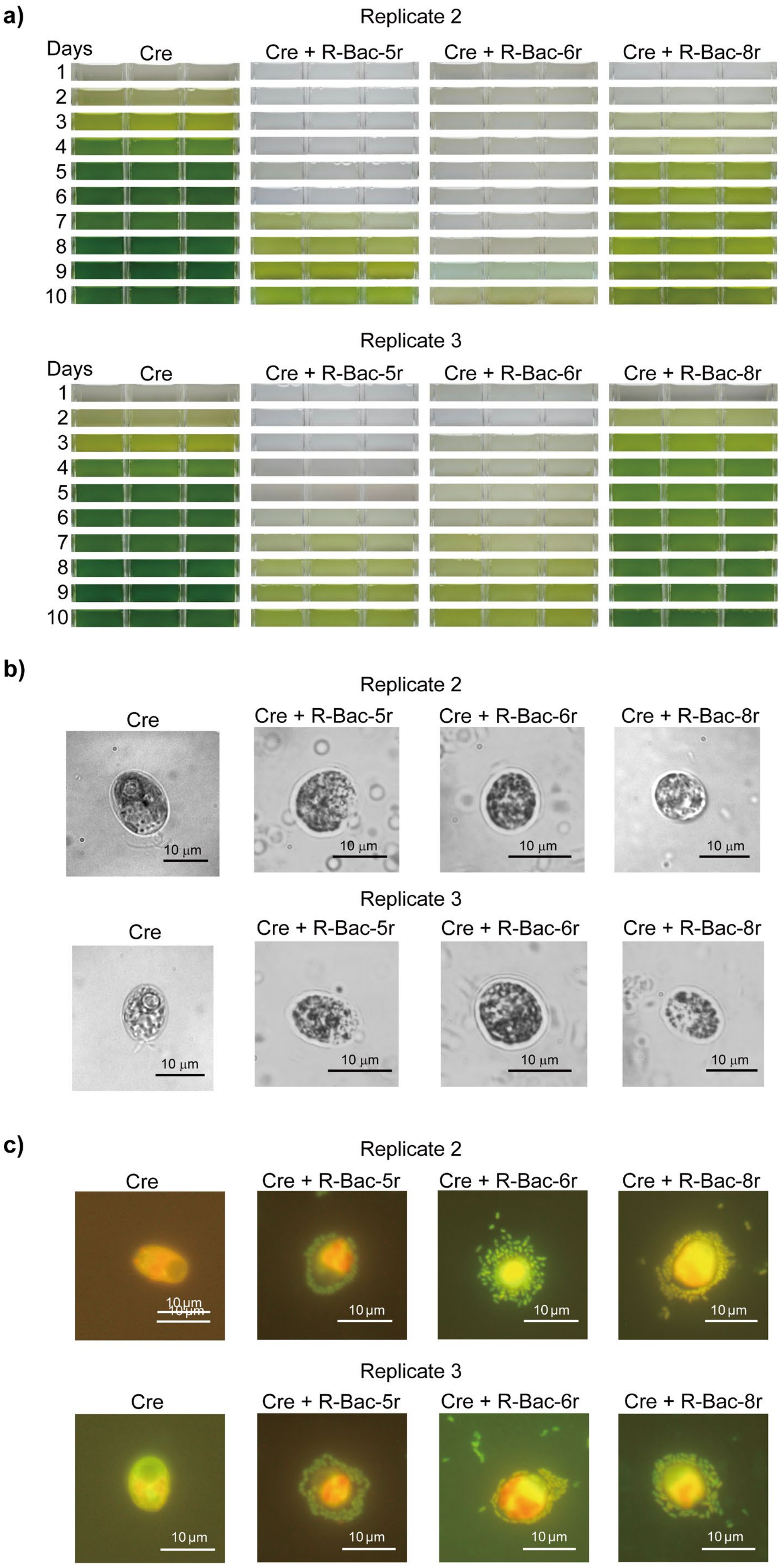
a)
Replicate 2
Days
Cre
Cre + R-Bac-5r
Cre + R-Bac-6r
Cre + R-Bac-8r
1
2
3
4
5
6
7
8
9
10
Replicate 3
Days
Cre
Cre + R-Bac-5r
Cre + R-Bac-6r
Cre + R-Bac-8r
1
2
3
4
5
6
7
8
9
10
b)
Replicate 2
Cre
Cre + R-Bac-5r
Cre + R-Bac-6r
Cre + R-Bac-8r
10 µm
Replicate 3
Cre
Cre + R-Bac-5r
Cre + R-Bac-6r
Cre + R-Bac-8r
10 µm
c)
Replicate 2
Cre
Cre + R-Bac-5r
Cre + R-Bac-6r
Cre + R-Bac-8r
10 µm
Replicate 3
Cre
Cre + R-Bac-5r
Cre + R-Bac-6r
Cre + R-Bac-8r
10 µm

**FIGURE S7 |** Cocultivation, cellular morphology and physical association of *C. reinhardtii* (Cre) with antagonistic *Pseudomonas* spp. (a-c) Experiments were performed in mono- and cocultures of Cre with the bacteria R-Bac-5r, R-Bac-6r, and R-Bac-8r, respectively, isolated from the rice rhizosphere. (a) Liquid mono- and cocultures of Cre and each of the above-mentioned antagonistic bacteria. The organisms were cocultivated in TAP medium (Harris 1989) at a ratio of 1:250 (algae:bacteria). Replicates 2 and 3 are presented. Replicate 1 is shown in Figure 1f. (b) Bright-field microscopy images of Cre cells in monoculture or cocultivated with R-Bac-5r, R-Bac-6r, and R-Bac-8r, respectively, for 24 h. Replicates 2 and 3 are shown; replicate 1 is shown in Figure 3a. (c) Acridine orange staining (Methods) visualizing the physical attachment of Cre cells and R-Bac-5r, R-Bac-6r, as well as R-Bac-8r after 10 min of incubation. Replicates 2 and 3 are shown; Replicate 1 is presented in Figure 3b.

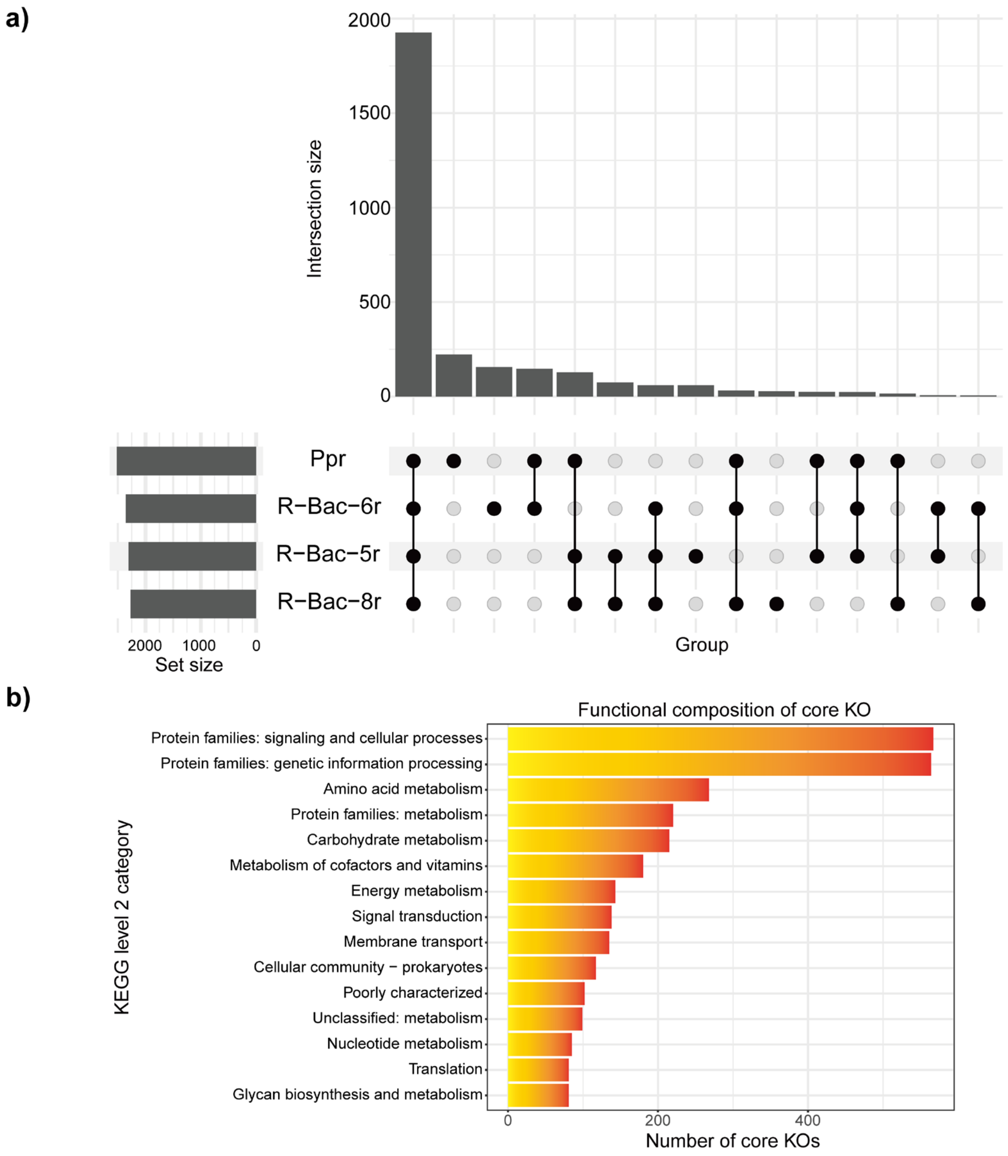


**FIGURE S8** | Comparative genomic analysis of *Pseudomonas protegens* (Ppr) and antagonistic rice rhizosphere isolates R-Bac-5r, R-Bac-6r, and R-Bac-8r. (a) UpSet plot showing the distribution and overlap of KEGG Orthology (KO) groups among Ppr and the three antagonistic isolates. In the UpSet plot, black filled circles indicate the presence of a KO group in the corresponding bacterial genome, whereas light gray circles indicate its absence (Lex et al. 2014). Black vertical lines connect the genomes containing the same KO group within each intersection, highlighting shared functional profiles among the genomes. The intersection bars above the matrix indicate the number of KO groups associated with each specific combination of genomes, while the set-size bars on the left represent the total number of KO groups identified in each genome. (b) Functional classification of core KO groups shared among the four bacterial genomes.

**Table S1** Overview of 12 bacterial isolates from either the rhizosphere (r) or the surrounding soil (s) from a rice field in Ottobiano, Italy.

| ID | Shape | Color | Opacity | Source |
|---|---|---|---|---|
| R-Bac-1s | Circular | Bright yellow | Opaque | Soil |
| R-Bac-2s | Circular | White | Opaque | Soil |
| R-Bac-3s | Circular | White | Opaque | Soil |
| R-Bac-4r | Circular | White to beige | Opaque | Rhizosphere |
| R-Bac-5r | Circular | Beige | Opaque | Rhizosphere |
| R-Bac-6r | Circular | Beige | Opaque | Rhizosphere |
| R-Bac-7s | Circular | Light yellow | Opaque | Soil |
| R-Bac-8r | Circular | White to beige | Opaque | Rhizosphere |
| R-Bac-9r | Circular | White | Opaque | Rhizosphere |
| R-Bac-10s | Circular | White | Opaque | Soil |
| R-Bac-11s | Circular | White | Translucent | Soil |
| R-Bac-12s | Circular | White | Opaque | Soil |

**Table S2** *16S rRNA* gene sequence dataset obtained from public databases.

| Number | Species | Source |
|---|---|---|
| 1 | *Paenarthrobacter histidinolovorans* A3 | GenBank: PQ856804.1 |
| 2 | *Enterobacter ludwigii* 22STR477 | GenBank: OR831825.1 |
| 3 | *Enterobacter sichuanensis* E2-10 | GenBank: OQ753127.1 |
| 4 | *Enterobacter asburiae* MN11SW4 | GenBank: JX501721.1 |
| 5 | *Enterobacter quasiroggenkampii* WCHECL1060 | GenBank: KY979139.1 |
| 6 | *Pseudomonas monteilii* JPR31 | GenBank: KR045595.1 |
| 7 | *Enterobacter asburiae* IHB B 1070 | GenBank: KF475833.1 |
| 8 | *Pseudomonas plecoglossicida* BOE7 | GenBank: MH580213.1 |
| 9 | *Pseudomonas putida* BGRI EBC SK23-T2 | GenBank: OR166199.1 |
| 10 | *Pseudomonas hunanensis* P11 | GenBank: MN946623.1 |
| 11 | *Aquipseudomonas alcaligenes* W5-8 | GenBank: MG905285.1 |
| 12 | *Aquipseudomonas alcaligenesm* AAUCP 3 | GenBank: MZ636709.1 |
| 13 | *Pseudomonas tohonis* Ca-76 | GenBank: OR781344.1 |
| 14 | *Pseudomonas aeruginosa* NR_114472 | GenBank: ON606274.1 |
| 15 | *Pseudomonas solani* Sm006$^{T}$ | GenBank: LC744517.1 |
| 16 | *Paenarthrobacter nicotinovorans* FLP55 | GenBank: MW131230.1 |
| 17 | *Pseudomonas shirazica* TNB-16 | GenBank: OL455721.1 |
| 18 | *Pseudomonas asiatica* N21 | GenBank: PV645863.1 |
| 19 | *Pseudomonas taiwanensis* BGRI EBC SS23-S7 | GenBank: OR166229.1 |
| 20 | *Pseudomonas entomophila* KLP 01 | GenBank: MW250866.1 |
| 21 | *Pseudomonas juntendi* AUMC B-549 | GenBank: PQ220143.1 |
| 22 | *Pseudomonas plecoglossicida* FC2949 | GenBank: MK089548.1 |
| 23 | *Pseudomonas alloputida* P4N15 | GenBank: OK668302.1 |
| 24 | *Pseudomonas putida* DSS-CAP-7 | GenBank: MF076564.1 |
| 25 | *Pseudomonas vancouverensis* R14 | GenBank: OM993328.1 |
| 26 | *Pseudomonas fluorescens* JB18 | GenBank: MZ156988.1 |
| 27 | *Pseudomonas koreensis* Ps 9-14 | NCBI Reference Sequence: NR_025228.1 |
| 28 | *Pseudomonas umsongensis* Ps 3-10 | NCBI Reference Sequence: NR_025227.1 |
| 29 | *Pseudomonas lutea* 6S1 | GenBank: MH179098.1 |
| 30 | *Pantoea agglomerans* DSM 3493$^{T}$ | NCBI Reference Sequence: NR_041978.1 |
| 31 | *Enterobacter huaxiensis* WTB145 | GenBank: MK734322.1 |
| 32 | *Pseudomonas paraglycinae* R45 | GenBank: LC807643.1 |
| 33 | *Pseudomonas moraviensis* 1B4$^{T}$ | NCBI Reference Sequence: NR_043314.1 |
| 34 | *Pseudomonas chlororaphis* WG405 | GenBank: PQ781434.1 |
| 35 | *Pseudomonas fitomaticsae* BGRI EBC SK23-T15 | GenBank: OR166212.1 |
| 36 | *Pseudomonas granadensis* 26C3 | GenBank: MG269614.1 |
| 37 | *Pseudomonas teessidea* B26 | GenBank: KT767690.1 |
| 38 | *Pseudomonas kribbensis* XG107 | GenBank: PQ781461.1 |
| 39 | *Pseudomonas protegens* CHA0 | NCBI Reference Sequence: NR_114749.1 |
| 40 | *Pseudomonas protegens* Pf-5 | GenBank: CP000076.1 |

**References**

Harris, E. H. 1989. *The Chlamydomonas sourcebook. A comprehensive guide to biology and laboratory use*. Academic Press.

Lex, A., Gehlenborg, N., Strobelt, H., Vuillemot, R., and Pfister, H. 2014. "UpSet: visualization of intersecting sets." *IEEE Transactions on Visualization and Computer Graphics* 20:1983–1992. https://doi.org/10.1109/TVCG.2014.2346248.
